# Genomic footprints of activated telomere maintenance mechanisms in cancer

**DOI:** 10.1101/157560

**Authors:** Lina Sieverling, Chen Hong, Sandra D. Koser, Philip Ginsbach, Kortine Kleinheinz, Barbara Hutter, Delia M. Braun, Isidro Cortés-Ciriano, Ruibin Xi, Rolf Kabbe, Peter J. Park, Roland Eils, Matthias Schlesner, Karsten Rippe, David T.W. Jones, Benedikt Brors, Lars Feuerbach, on behalf of the PCAWG Structural Variation Working Group, PCAWG SNV Calling Working Group, PCAWG Drivers and Functional Interpretation Group, PCAWG Evolution and Heterogeneity Working Group, PCAWG Technical Working Group, and the ICGC/TCGA Pan-Cancer Analysis of Whole Genomes Network

## Abstract

Cancers require telomere maintenance mechanisms for unlimited replicative potential. We dissected whole-genome sequencing data of over 2,500 matched tumor-control samples from 36 different tumor types to characterize the genomic footprints of these mechanisms. While the telomere content of tumors with *ATRX* or *DAXX* mutations (ATRX/DAXX^trunc^) was increased, tumors with *TERT* modifications showed a moderate decrease of telomere content. One quarter of all tumor samples contained somatic integrations of telomeric sequences into non-telomeric DNA. With 80% prevalence, ATRX/DAXX^trunc^ tumors display a 3-fold enrichment of telomere insertions. A systematic analysis of telomere composition identified aberrant telomere variant repeat (TVR) distribution as a genomic marker of ATRX/DAXX^trunc^ tumors. In this clinically relevant subgroup, singleton TTCGGG and TTTGGG TVRs (previously undescribed) were significantly enriched or depleted, respectively. Overall, our findings provide new insight into the recurrent genomic alterations that are associated with the establishment of different telomere maintenance mechanisms in cancer.

---

Telomeres are nucleoprotein complexes at the ends of chromosomes that prevent DNA degradation and genome instability^1^. The typically 10-15 kb long chromosome termini are composed of long stretches of TTAGGG (t-type) repeat arrays with an increasing number of variants towards proximal, subtelomeric regions, the most common being TGAGGG (g-type), TCAGGG (c-type) and TTGGGG (j-type) repeats^2,3^.

Telomeres play an important role in cellular aging, as they are shortened with each cell division and finally trigger a DNA-damage response resulting in senescence^4,5^. To avoid this permanent growth arrest, cells with unlimited proliferative potential need to extend their telomeres. In humans, telomeric DNA is synthesized onto the chromosome ends by telomerase, an enzyme that is composed of the reverse transcriptase TERT and the RNA template TERC. This complex is active in the germline and stem cells, but absent in most somatic cells^6^. Telomerase is up-regulated in about 85% of human cancers by different mechanisms, including *TERT* amplifications^7^, rearrangements^8-10^ or mutations in the *TERT* promoter^11,12^. The remaining tumors employ an alternative lengthening of telomeres (ALT) pathway, which is based on DNA recombination of telomeric sequences^13^. Details on the ALT mechanism remain elusive but it has been associated with loss-of-function mutations in the chromatin remodeling genes *ATRX* (α-thalassaemia/mental retardation syndrome X-linked) and *DAXX* (death-domain associated protein)^14^. Telomeres of ALT cells characteristically have heterogeneous lengths and contain a range of telomere variant repeats (TVRs)^15-17^. Other hallmarks of ALT include ALT-associated promyelocytic leukemia (PML) nuclear bodies (APBs), abundance of extrachromosomal telomeric repeats of various forms (such as C-circles) and genome instability^13,18^.

While normally located at the chromosome termini, telomere sequences are also found in intrachromosomal regions. As such, interstitial telomeric sequences with large blocks of telomere repeats exist in humans and other species, which probably arose from ancestral genome rearrangements or other evolutionary events^19^. Recently, ALT-specific, targeted telomere insertions into chromosomes that lead to genomic instability have also been described^20^. Another source for unexpected telomere repeat sites is the stabilizing function of telomeres at broken chromosomes. After a double-strand break, telomeres can be added *de novo* to the unprotected break sites (“telomere healing”)^21,22^ or acquired from other chromosomal positions (“telomere capture”)^23,24^.

Here, we characterized the telomere landscape of 2,519 tumor samples from 36 different tumor types using whole genome sequencing data from the Pan-Cancer Analysis of Whole Genomes (PCAWG) 25 project. Besides determining telomere content and searching for mutations associated with different telomere maintenance mechanisms (TMMs), we systematically detected 2,683 somatic telomere insertions and show that different TMMs are associated with enrichment of previously undescribed singleton TVRs.

## Results

### Telomere content across cohorts

Due to the repetitive nature of telomere sequences, short sequencing reads from telomeres cannot be uniquely aligned to individual chromosomes. However, a mean telomere content for the tumor as a whole can be estimated from the number of reads containing telomere sequences^17,26-29^. Here, we extracted reads containing at least six telomere repeats per 100 bases, allowing the canonical telomere repeat TTAGGG and the three most common TVRs TCAGGG, TGAGGG and TTGGGG. The telomere content was defined as the number of unaligned telomere reads normalized by sequencing coverage and GC-content. Of the 2,583 high-quality tumor samples available in PCAWG, we selected those from donors with a single tumor sample. The telomere content was determined for the remaining 2,519 tumor samples and matched controls from 36 different tumor types. All relevant donor information and results used in this study are summarized in Supplementary Table 1.

Telomere content of the controls anti-correlated with age (r = -0.36, Spearman correlation) (Supplementary Fig. 1a). However, this age effect only has a low contribution to the strong correlation between the telomere content of the tumor and control samples (r = 0.47 and r_partial_ = 0.46 given the patient age, Spearman correlation, Supplementary Fig. 1b). Thus, the association of tumor and control telomere content must mainly be caused by other genetic^30,31^, environmental^32^ or technical factors^33^. We normalized for this by computing the ratio of tumor and control telomere content per individual.

Most tumor samples had a lower telomere content than the matched control (Fig. 1a). However, there were systematic differences between the different tumor types. Among those with the highest telomere content increase were osteosarcomas and leiomyosarcomas (median telomere content tumor/control log2 ratios = 0.7 and 0.6, respectively). A particularly low telomere content was found in colorectal adenocarcinoma and medulloblastoma (median telomere content tumor/control log2 ratios = -1.0).

**Figure 1:**
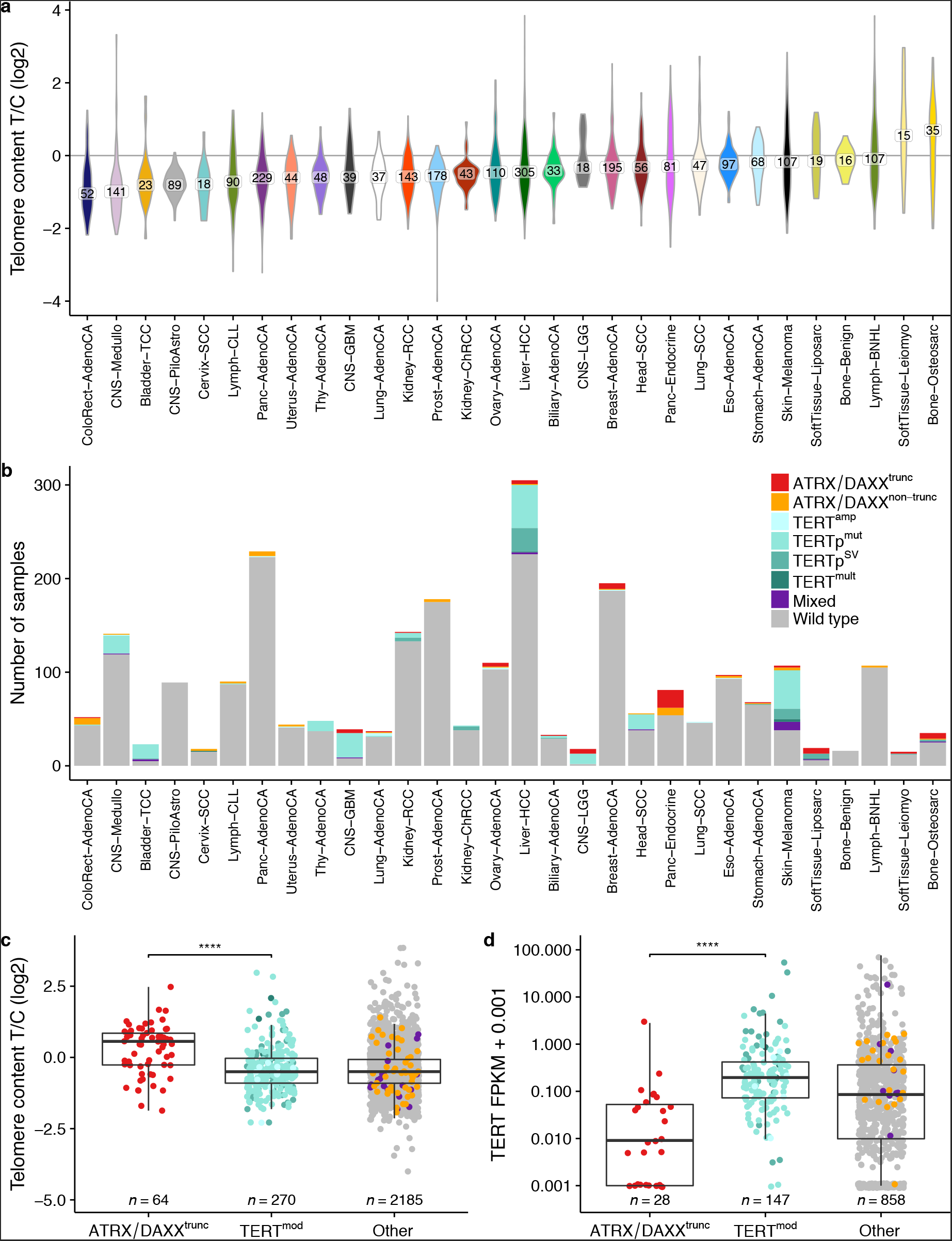
Telomere content is increased in ATRX/DAXX^trunc^ samples. (a) Overview of the telomere content distribution of all analyzed tumor types. The number of samples in each tumor type is indicated. Cohorts with sample sizes below 15 are not shown. (b) TMM-associated mutations in different tumor types. (c) Telomere content in samples with different TMM-associated mutations. (d) *TERT* expression in samples with different TMM-associated mutations. \*\*\*\**p* < 0.0001, Wilcoxon rank-sum tests.

### Prevalence of TMM-associated mutations

Different types of mutations in *ATRX* or *DAXX* and at the *TERT* locus have been associated with ALT and telomerase activation, respectively. We therefore searched for these types of somatic mutations to infer the active TMM in a given tumor. Somatic mutations in *ATRX*, *DAXX* or *TERT* were found in 16% of tumor samples. In total, 64 tumor samples had truncating *ATRX* (*n* = 53) or *DAXX* alterations (*n* = 11) and are referred to as *ATRX/DAXX*^trunc^ in the following analysis. Of note, 10 of the 11 *DAXX* alterations were found in pancreatic endocrine tumors, while *ATRX* mutations were seen in a wider variety of entities. An additional 46 samples had non-truncating *ATRX/DAXX* simple nucleotide variants. *TERT* alterations were detected in 270 tumor samples (TERT^mod^). The latter group comprised 198 activating C228T or C250T promoter mutations (of which 132 were obtained from the PCAWG simple nucleotide variant consensus calls and the remaining were detected with a less stringent approach), 11 amplifications leading to at least six additional *TERT* copies, 55 structural variations within 20 kb upstream of *TERT*, and 6 samples with more than one of these modifications. Additionally, 18 tumor samples had both *ATRX/DAXX* truncating or other missense mutations and *TERT* alterations.

The structural variations upstream of *TERT* showed a strikingly focal distribution close to the transcription start site (TSS) (Supplementary Fig. 2), suggesting an advantage of tumors with such rearrangements. Moreover, 40% (*n* = 25/62) of the juxtaposed positions within 20 kb upstream of the *TERT* TSS overlapped directly with enhancers from the dbSUPER database^34^. In contrast, only 13% (n = 9/69) of the juxtaposed positions between 20 and 1,000 kb corresponded to a predicted superenhancer. These results point to “enhancer hijacking” near the *TERT* TSS, a phenomenon which has been described in neuroblastoma^8,9^ and for which indications have recently been found in further cancer types^33^.

The tumor types with the highest prevalence of ATRX/DAXX^trunc^ mutations were liposarcomas (32%), adult lower grade gliomas (28%), pancreatic endocrine tumors (23%) and osteosarcoma (17%) (Fig. 1b), all of which have previously been associated with ALT^14,35^. Importantly, the five lower grade glioma samples with *ATRX* alterations are not oligodendrogliomas according to the recent WHO classification^36^, as they are lacking 1p/19q co-deletions. TERT^mod^ were most prevalent in transitional cell bladder cancer (70%), glioblastoma (67%), lower grade gliomas (61%) and melanoma (51%).

The telomere content in TERT^mod^ samples differed significantly from that in ATRX/DAXX^trunc^ samples (*p* = 1.1 × 10^−9^, Wilcoxon rank-sum test; Fig. 1c, a detailed overview is shown in Supplementary Fig. 3a). On average, telomere content was gained in ATRX/DAXX^trunc^ (mean telomere content tumor/control log2 ratio = 0.3), while telomere sequences were lost in TERT^mod^ samples (mean telomere content tumor/control log2 ratio = -0.4). Samples with non-truncating *ATRX/DAXX* simple nucleotide variants had a similar telomere content as TERT^mod^ samples (*p* > 0.05, Wilcoxon rank-sum test), suggesting that most of the non-truncating *ATRX/DAXX* mutations are passenger events. In TERT^mod^ samples and samples with unknown TMM, the telomere content correlated with *TERT* expression (r = 0.20, Pearson -10 correlation; *p* = 4.1 × 10^-10^, significance of fitted linear regression model) and *TERT* expression was significantly higher in TERT^mod^ samples than in ATRX/DAXX^trunc^ samples (*p* = 1.3 × 10^−9^, Wilcoxon rank-sum test; Fig. 1d, a detailed overview is shown in Supplementary Fig. 3b).

### Telomere insertions occur frequently in tumors with ALT-associated mutations

To find insertions of telomeres into non-telomeric regions of the genome, we searched for tumor-specific discordant paired-end reads where one end maps to the chromosome and the other end is telomeric. Exact positions of the insertions were determined from reads spanning the junction site and visually inspected (examples in Fig. 2).

Overall, 2,683 telomere insertions were detected. These were distributed unevenly between samples and different tumor types (Fig. 3a). Telomere insertions were found in 27% of the tumor samples, with counts ranging between one and 228 telomere insertion events. The tumor types with the highest amount of telomere insertions per tumor sample were liposarcoma, leiomyosarcoma and osteosarcoma, all of which also had a relatively high mean telomere content. In fact, the number of telomere insertions positively correlated with the telomere content (r = 0.19, Spearman correlation). Moreover, the number of telomere insertions was associated with the number of genomic break points in the sample (r = 0.38, Spearman correlation). To test for a synergistic effect, linear models that predict telomere insertions from telomere content and breakpoint abundance with and without an interaction term were computed. -234 The models with the interaction term (*p* = 8.8 × 10) performed substantially better than purely additive models (*p* = 5.8 × 10^−90^).

There was clearly a higher percentage of samples with telomere insertions in ATRX/DAXX^trunc^ tumors (80%) than TERT^mod^ tumors (28%) (Fig. 3b). As expected, ATRX/DAXX^trunc^ samples also had a higher number of breakpoints (mean = 733) than TERT^mod^ samples (mean = 291) (Fig. 3c). In keeping with this, chromothripsis (numerous chromosomal rearrangements occurring in a single event)^37^ was more prevalent in the ATRX/DAXX^trunc^ samples (59%) compared to TERT^mod^ samples (34%) and samples without ATRX/DAXX^trunc^ and TERT^mod^ mutations (29%). Overall, the fraction of genomic breakpoints overlapping with telomere insertion sites was significantly higher in ATRX/DAXX^trunc^ than TERT^mod^ -20 samples (*p* = 1.7 × 10^-20^, Wilcoxon rank-sum test; Fig. 3d). Correlation analysis of telomere insertions and mutations in telomere maintenance-associated genes from the TelNet database^38^ (http://www.cancertelsys.org/telnet) revealed significant association with *TP53* (*q* = 1.9 × 10^−42^), *ATRX* (*q* = 2.6 × 10^−6^), *PLCB2* (q = 7.8 × 10^−4^), *MEN1* (*q* = 0.017), *TSSC4* (*q* = 0.017), *RB1* (*q* = 0.018), *DAXX* (*q* = 0.019) and *ABCC8* mutations (*q* = 0.04, Wilcoxon rank-sum tests after Benjamini-Hochberg correction). Most of these genes have been implicated in the maintenance of telomere length or structure in humans (Supplementary Table 2). The exceptions are *PLCB2* and *ABCC8*, whose homologues have so far only been reported in association with telomere length regulation in yeast^39,40^.

The detected telomere insertions were scattered across different chromosomes and regions within the chromosome (Supplementary Fig. 4). No clear preferential insertion sites were identified, but several *de novo* telomere junctions occurred at the chromosome ends (5% within 50 kb of the first or last chromosomal segment). A total of 44% of the telomere insertions were in genes, and 8% of these disrupted exons. Several tumor suppressor genes were affected, e.g. *CHEK1* encoding for a protein involved in cell cycle arrest upon DNA damage^41^ (Fig. 2a).

Of note, patterns of microhomology were observed in 79% of telomere insertions with t-type repeats at the junction site (Supplementary Fig. 5).

**Figure 2:**
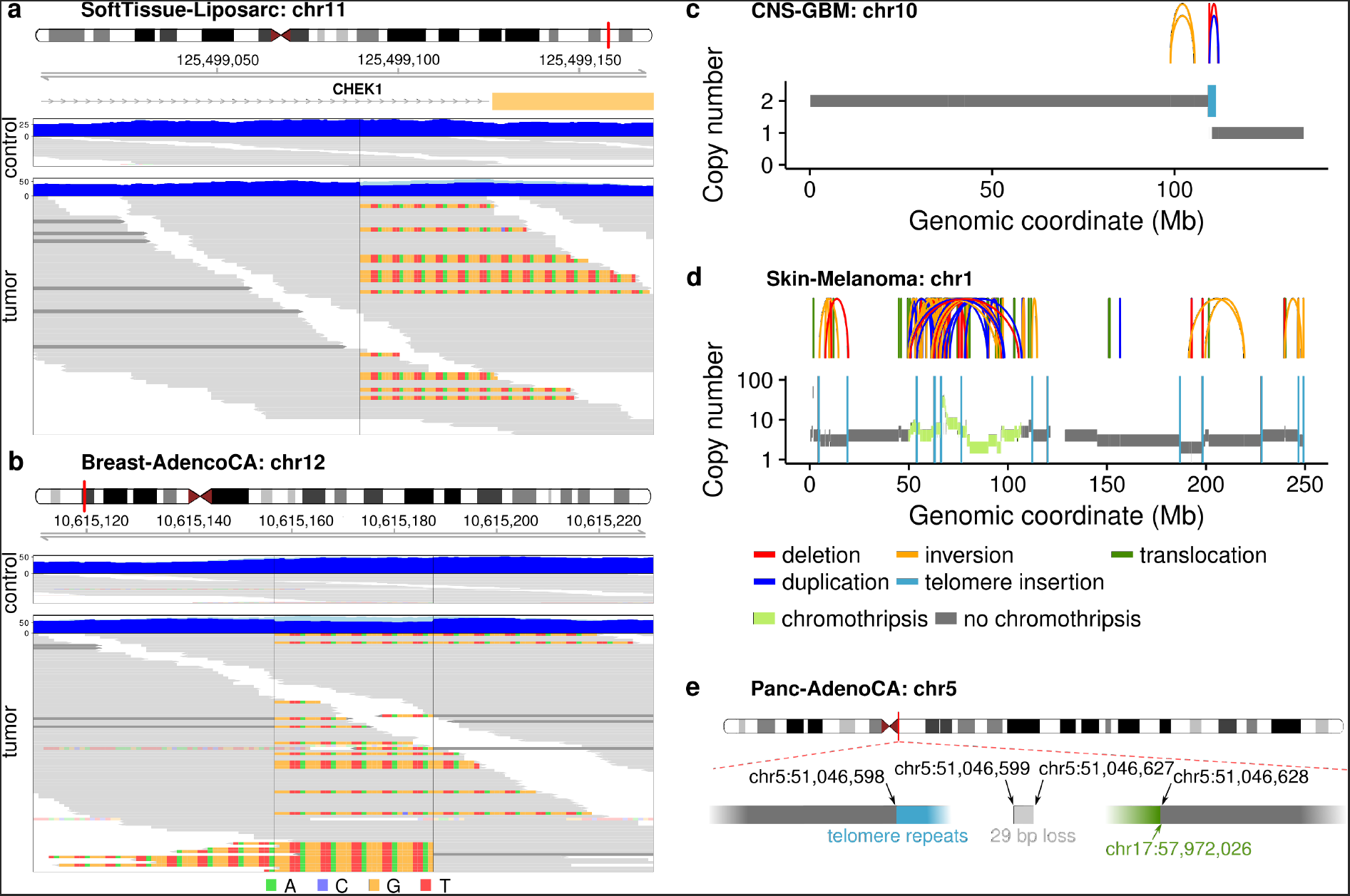
Examples of telomere insertions. (a) One-sided telomere insertion in liposarcoma sample SP121774. Blue tracks show the sequencing coverage; light blue represents clipped sequences. Individual reads are grey and clipped bases are colored. Dark grey reads represent the non-telomeric end of a discordant read pair. (b) Two-sided telomere insertion in breast adenocarcinoma sample SP5636. Non-telomeric clipped bases are transparent. (c) One-sided telomere insertion accompanied by copy number loss of the adjacent chromosome end in glioblastoma sample SP29559. Arches represent structural variations. (d) Multiple telomere insertions in chromosome that underwent chromothripsis in melanoma sample SP124441. (e) One-sided telomere insertion accompanied by a translocation of the adjacent chromosome segment in pancreatic adenocarcinoma sample SP125764.

**Figure 3:**
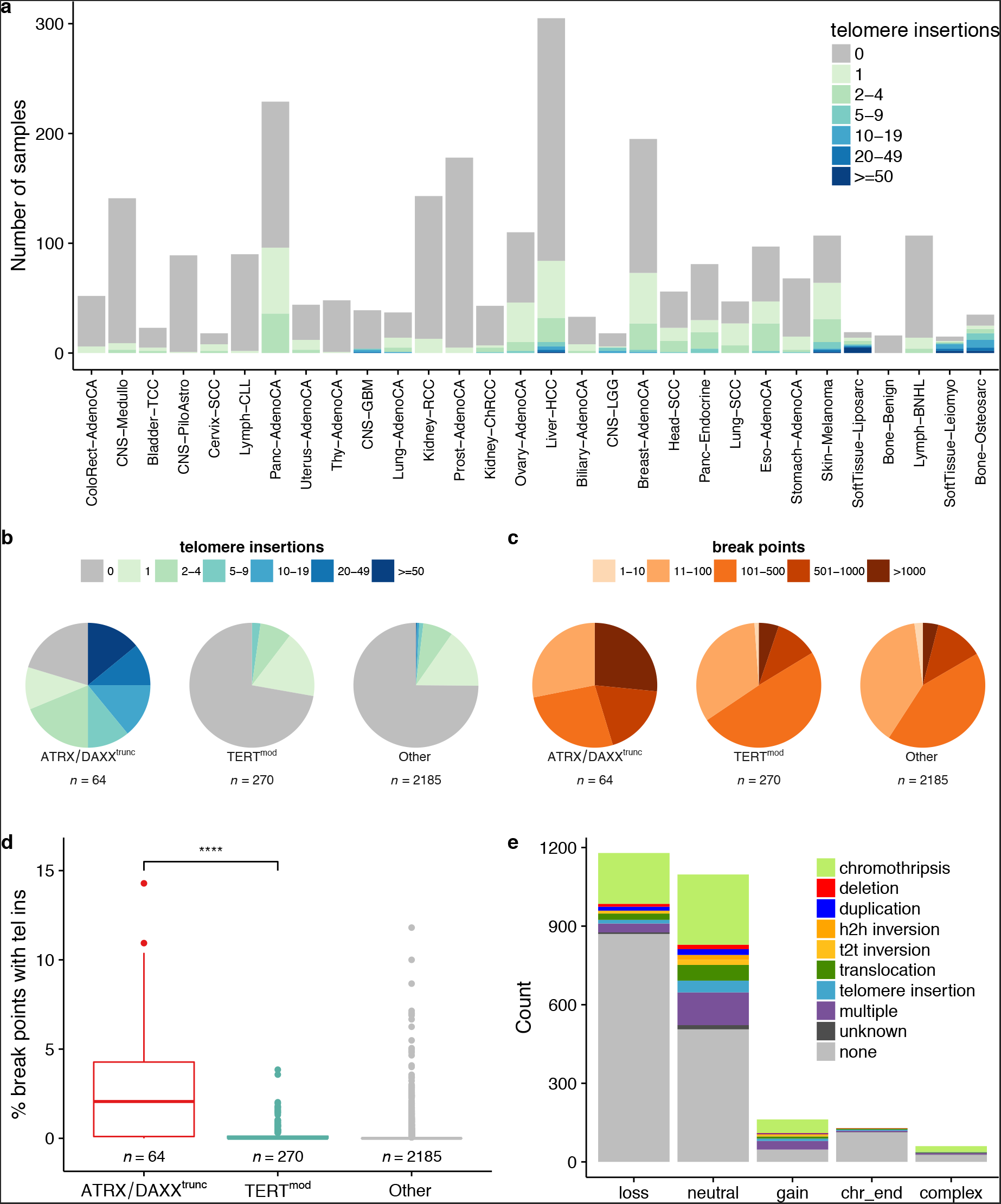
Insertion of telomere sequences into non-telomeric chromosomal regions. (a) Number of telomere insertions in samples of different tumor types. The tumor types are sorted by mean telomere content tumor/control log2 ratios. Cohorts with sample sizes below 15 are not shown. (b) Number of telomere insertions in samples with different TMM-associated mutations. (c) Number of break points in samples with different TMM-associated mutations. (d) Percent of break points coinciding with telomere insertions in samples with different TMM-associated mutations. \*\*\*\**p* < 0.0001, Wilcoxon rank-sum test. (e) Copy number changes of adjacent segments accompanying telomere insertions. “Complex” means that the copy numbers between segments differ in more than four copies. Overlaps with regions of chromothripsis are indicated. For telomere insertions that did not overlap with regions of chromothripsis, structural variations or additional telomere insertions within 10 kb are indicated. h2h = head-to-head, t2t = tail-to-tail.

### Telomere insertions often coincide with loss of the adjacent chromosomal segment

Most of the telomere insertions were one-sided (98%), i.e. telomere sequences were only attached to one side of the breakpoint (Fig. 2a). Telomere insertions were defined as two-sided if there was a second telomere insertion event downstream in the opposite orientation (Fig. 2b). Because so many breakpoints were one-sided, we investigated the fate of the corresponding broken fragment using complementary information from copy number changes and structural variation annotation (Fig. 3e). As expected, onesided telomere insertions coincided most frequently with copy number loss of the adjacent segment (46%, Fig. 2c). In contrast, copy number gains of the fragment were rare (6%). Surprisingly, telomere insertions were frequently located at copy number neutral sites (42%). Overlaps with regions of chromothripsis were found for 25% and structural variations without chromothripsis overlap (including telomere insertions) were detected near the insertion site for 28% of the copy-number neutral cases (Fig. 2d-e). The remaining telomere insertions at copy-number neutral sites are likely to be subclonal (Supplementary Fig. 6a) or have undetected structural variations nearby (Supplementary Fig. 6b).

### Singleton TVRs are enriched in ATRX/DAXX^trunc^ samples

It has previously been shown that ALT leads to an increased integration of TVRs into telomeres, the most common ones being hexamers of the type NNNGGG^17^. To detect differences in the telomere composition of ATRX/DAXX^trunc^ and TERT^mod^ tumors, we therefore searched for NNNGGG repeats in telomere reads. The most frequent TVRs across all tumor samples were TGAGGG, TCAGGG and TTGGGG (Supplementary Fig. 7), which are known to be enriched in proximal telomeric regions^2,3^.

These and the seven other most frequent TVRs (TAAGGG, GTAGGG, CATGGG, TTCGGG, CTAGGG, TTTGGG and ATAGGG) were chosen to search for common telomere repeat combinations. For this, the neighboring 18 base pairs on either side of the TVRs were determined (Supplementary Table 3). Most TVRs were surrounded by many different pattern combinations (e.g. TTGGGG). Others were dominated by a certain repeat context, which was similar in ATRX/DAXX^trunc^ and TERT^mod^ tumors (e.g. CATGGG or ATAGGG). However, TTCGGG stood out, as 35% of the TVRs in ATRX/DAXX^trunc^ samples were surrounded by canonical t-type repeats, whereas this context was observed for only 2% of TTCGGG TVRs in TERT^mod^ tumors.

Following up on this observation, we compared variant hexamers surrounded by at least three t-type repeats on either side (“singletons”) to TVRs in an arbitrary sequence context. This revealed that singletons are generally well suited to distinguish ATRX/DAXX^trunc^ from TERT^mod^ samples (Fig. 4a, an overview of all patterns is shown in Supplementary Fig. 8). The remaining variant analysis therefore focused on such TVR singletons. CATGGG was excluded as it did not occur as singletons. For the other TVRs, the median of absolute counts varied between 13 and 112, but counts in individual tumor samples reached more than ten thousand (Supplementary Fig. 9).

As expected, normalized singleton repeat counts generally rose with increasing telomere content (Fig. 4b, an overview of all patterns is shown in Supplementary Fig. 10). However, TGAGGG, TCAGGG, TTGGGG and TTCGGG singletons had significantly higher counts than expected in ATRX/DAXX^trunc^ compared to TERT^mod^ samples (*p* = 8.2 × 10^−9^, 2.3 × 10^−5^, 4.4 × 10^−4^, and 3.2 × 10^−12^, respectively, Wilcoxon rank sum test after Bonferroni correction; Fig. 4c). Especially TGAGGG and TTCGGG seemed to be highly interspersed in a subset of ATRX/DAXX^trunc^ tumors. In contrast, TTTGGG singletons were observed less frequently in ATRX/DAXX^trunc^ tumors (*p* = 3.6 × 10^−12^, Wilcoxon rank sum test after Bonferroni correction).

This seemingly ALT-specific TVR enrichment or depletion occurred in different tumor types, with the highest prevalence in leiomyosarcoma (60%), pancreatic endocrine tumors (42%), osteosarcomas (29%) and lower grade gliomas (28%; Supplementary Table 4). In the ATRX/DAXX^trunc^ samples, singleton TVR occurrences correlated with each other (Supplementary Fig. 11). The strongest correlations were between TGAGGG occurrence and TTCGGG and TTGGGG singletons (both r = 0.56, Spearman correlation).

**Figure 4:**
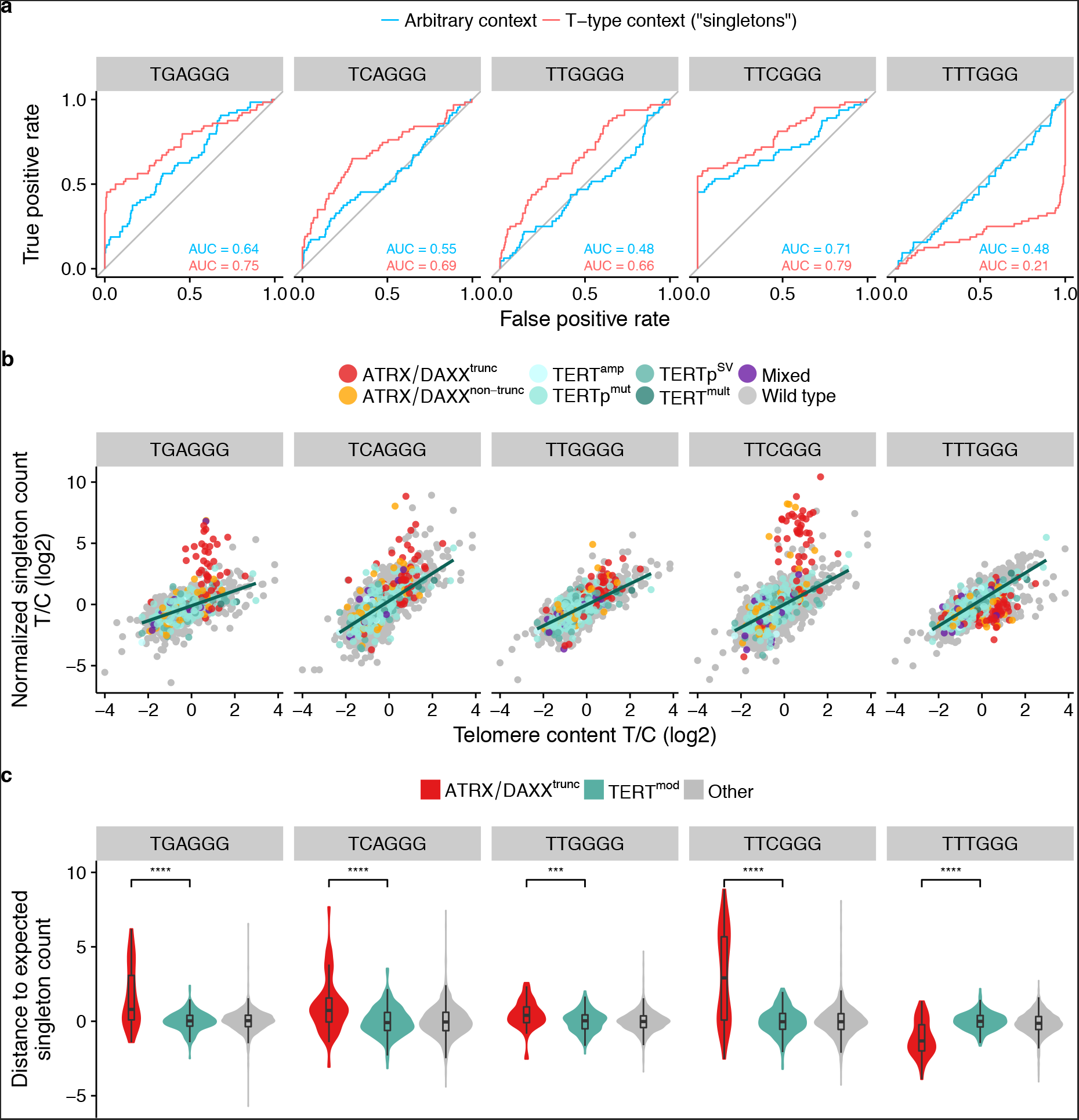
Singleton TVRs enriched or depleted in ATRX/DAXX^trunc^ samples. (a) Receiver operating characteristic for the classification of samples with ALT-associated mutations from telomere variant repeats. Red: no specific sequence context required. Blue: singletons ((TTAGGG)3-NNNGGG-(TTAGGG)_3_). The more the area under the curve (AUC) deviates from 0.5, the better the repeat occurrence distinguishes ATRX/DAXX^trunc^ from TERT^mod^ samples. (b) Pattern count tumor/control log2 ratios of all patients plotted against telomere content tumor/control log2 ratios for selected singletons. The regression line through the TERT^mod^ samples is shown in green and is defined as the expected pattern count in the following. (c) Distance to the expected singleton repeat count in ATRX/DAXX^trunc^ and TERT^mod^ samples. \*\*\*\**p* < 0.0001; \*\**p* < 0.01, Wilcoxon rank-sum tests after Bonferroni correction. The profiles of all analyzed patterns are shown in Supplementary Figure 9 and 10.

### ALT prediction

ALT has several different hallmarks with which it can be reliably identified. However, none of these are detectable in short-read whole-genome sequencing data. Using ATRX/DAXX^trunc^ as indicators of ALT, we have shown several possible TMM classification features in this type of data.

Most ATRX/DAXX^trunc^ samples are already separated well from TERT^mod^ samples by non-supervised clustering of normalized TGAGGG, TCAGGG, TTGGGG, TTCGGG and TTTGGG singleton repeat counts (Supplementary Fig. 12). As expected, the clusters of ATRX/DAXX^trunc^ samples had a high telomere content and a high number of telomere insertions relative to the total number of breakpoints.

They were further used to build a random forest classifier distinguishing ATRX/DAXX^trunc^ from TERT^mod^ samples (area under the curve: 0.96; sensitivity: 0.72; specificity: 0.98; all after 10-fold cross-validation, Supplementary Fig. 13 and 14). The variables with the highest importance for the classification were the divergence of observed TTTGGG and TTCGGG singleton TVRs from the expected count, the number of breakpoints and the number of telomere insertions (Supplementary Table 5). It may be pivotal for further understanding of this mechanism to determine the causal relationship between these features and the ALT phenotype.

## Discussion

In this study, we have shown that the presence of ALT-associated mutations in tumors correlates with increased telomere content, enrichment of isolated TVRs in t-type context (singletons), a higher number of genomic breakpoints and intrachromosomal telomere insertions (Fig. 5). In contrast, tumors with mutations associated with a possible telomerase-activation showed moderate decrease of telomere content and increased *TERT* expression. Hence, *TERT* reactivation may a) not suffice to fully counteract the telomere loss associated with high proliferation and/or b) occur in later stages of tumor development when the telomeres have already reached a relatively short length. The observed telomere content increase in ATRX/DAXX^trunc^ versus the decrease in TERT^mod^ samples is in agreement with the recent findings of Barthel *et al.*^33^. The higher telomere content in ATRX/DAXX^trunc^ tumors indicates that the negative feedback loop that constrains telomere elongation to a physiological level in healthy telomerase-expressing cells^42,43^ is bypassed by the ALT process, while it seems to remain intact in telomerase-positive tumors. In addition to telomere elongation, the increase of telomere content in ALT-positive tumors detected by sequencing-based methods may partly stem from aberrant intrachromosomal telomere insertions^20^ or extra-chromosomal telomeric DNA^44^. Although almost all tumors must maintain their telomeres^45^, we only detected somatic mutations highly associated with ALT or telomerase activation in a subset of the samples. Most tumors must therefore activate telomere maintenance by other mechanisms, e.g. epigenetic control of *TERT* expression^33,46^.

**Figure 5:**
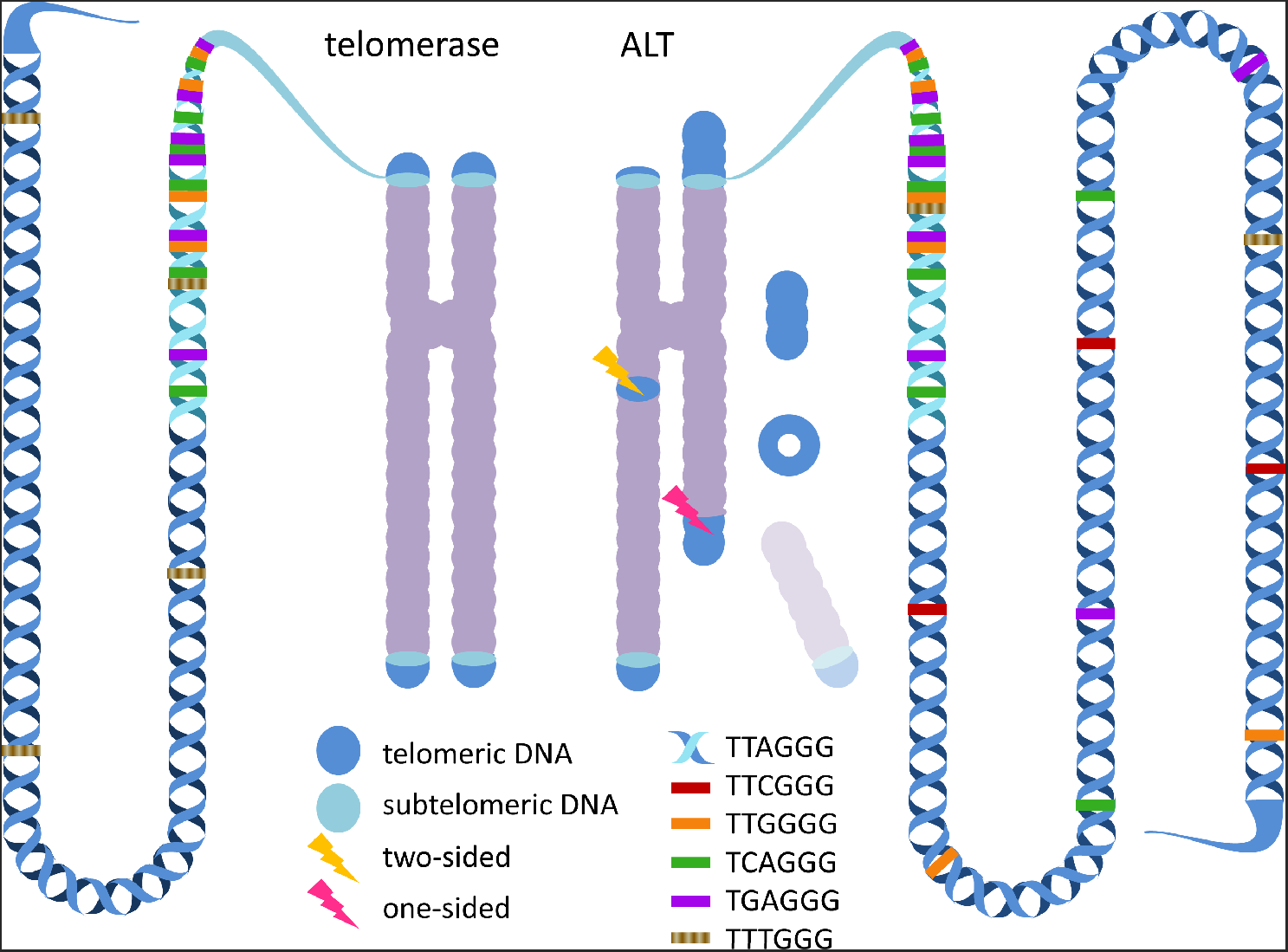
Genomic footprints of telomerase-mediated telomere elongation and ALT. It is known that telomeres elongated by telomerase have a homologous length with few TVRs in distal telomeric regions (left), while ALT telomeres have heterogeneous lengths with an increased amount of TVRs (right). Moreover, ALT cells have abundant extrachromosomal telomeric sequences. From this study, we conclude that the chromosomes of ALT cells have a higher number of aberrant interstitial telomere insertions, most of which are one-sided and accompanied by a loss of the adjacent chromosomal segment. We also showed that several TVRs occurring as singletons are more abundant in ALT telomeres, while one singleton (TTTGGG) was more abundant in telomerase-elongated telomeres. Please note that it is currently undetermined whether the different types of singletons are located in proximal or distal telomeric regions.

In our study, we systematically mapped telomere insertions into non-telomeric genomic regions using whole-genome sequencing data. They were most frequently accompanied by a loss of the adjacent chromosomal segment or located at copy number neutral sites. Surprisingly, the latter telomere insertions were rarely two-sided and chromothripsis or other structural variations in the adjacent genomic regions occurred only in about half of the cases. As broken chromosome ends are highly unstable, the remaining segments must have undetected structural rearrangements such as subclonal copy number changes or undetected DNA fusions. Taken together, the results suggest that we observe telomere healing or capture^22,23^ rather than telomere insertions followed by chromosomal instabilities^20,47^. As microhomology around telomere insertion sites was frequent, the sequences were probably inserted by nonhomologous end-joining^48^ or a microhomology-mediated mechanism^49^.

Telomere insertions were particularly frequent in ATRX/DAXX^trunc^ tumors, in which the abundant extrachromosomal telomeric DNA expands the telomere template pool for microhomology-mediated double strand repair. We speculate that in this cellular environment, a high load of genomic breakpoints subsequently leads to the observed disproportionately increased number of telomere capture-like events. Due to the stochastic nature of ALT, the likelihood of telomere crisis is elevated. The recently described induction of chromothripsis by telomere crisis^50^ may thus explain the observed higher prevalence of chromothripsis in ATRX/DAXX^trunc^ cases in this study.

Telomere elongation by ALT or telomerase enriches distinct TVRs^17^. Here, we report a stronger association of singleton TVRs with ATRX/DAXX^trunc^ mutations than TVRs in an arbitrary context. The increase of TVRs has been attributed to the inclusion of subtelomeric regions during ALT via homologous recombination^16^. Whether telomeric sequences with lower TVR density are under positive selection or regions with higher TVR density are under negative selection remains to be clarified.

A possible function for TVRs has been reported in ALT-positive cell lines, where TCAGGG repeats that recruit nuclear receptors were enriched^16,20^. This enrichment was confirmed in a subset of primary ATRX/DAXX^trunc^ tumor samples in our study, but was not as strong as the enrichment of TTCGGG or TGAGGG. While TGAGGG has previously been associated with ALT^16^, the high prevalence of TTCGGG singletons in ALT is a novel discovery. No proteins with strong affinity to these two TVRs are currently known. This may indicate a more passive mode of action, for instance deprotection of telomeres by shelterin displacement^16^, and/or alteration of the telomeric G-quadruplex conformation^17^. Notably, we report for the first time a hexamer which is depleted in ATRX/DAXX^trunc^ samples. More specifically, only TTTGGG singletons and not TTTGGG in arbitrary context show this pattern, which underlines the necessity to consider the sequence context of TVRs. None of the current models of ALT provide an explanation for this specific TVR depletion.

The here presented methodologies expand the established telomere content estimation from genomic sequencing by the context-dependent analysis of TVRs and telomere insertions, as well as in the scale of their application to a large pan-cancer study, thus adding new dimensions for the characterization of different telomere maintenance mechanisms.

## Methods

### Sequencing data

Whole genome sequencing (WGS) and expression data were obtained from the Pan-Cancer Analysis of Whole Genomes (PCAWG) project^25^. The WGS reads of tumor and control samples were aligned with bwa-mem by the PCAWG-tech group. Tumors with multiple samples were excluded from this study, as well as one sample pair with reads shorter than 30 bp. Expression data was in the format of normalized RNA read counts per gene and only available for 1,033 of 2,519 patients. Of note, we used the name “CNS-LGG” for the “CNS-Oligo” tumor type, because several samples in this cohort did not have the genetic markers for oligodendroglioma required by the WHO^36^.

### Mutation data

Somatic simple nucleotide, structural variations and copy numbers were obtained from the PCAWG consensus calls (Synapse IDs syn7364923, syn7596712 and syn8042992, respectively). Structural variations were not available for 24 tumor samples.

### Telomere read extraction and computational telomere content estimation

The telomere content of WGS samples was determined using the software tool TelomereHunter (www.dkfz.de/en/applied-bioinformatics/telomerehunter/telomerehunter.html)^51^. In short, telomeric reads containing six non-consecutive instances of the four most common telomeric repeat types (TTAGGG, TCAGGG, TGAGGG and TTGGGG) were extracted. For the further analysis, only unmapped reads or reads with a very low alignment confidence (mapping quality lower than 8) were considered. The telomere content was determined by normalizing the telomere read count by all reads in the sample with a GC-content of 48-52%.

### Determining TMM-associated mutations

Samples with a truncating *ATRX* or *DAXX* alteration (frame-shift insertion/deletion, stop-codon gain or structural variation breakpoint within the gene) were defined as ATRX/DAXX^trunc^, samples with other simple nucleotide variants were defined as ATRX/DAXX^non-trunc^. Deletions that only affected intronic regions of *ATRX* were not considered. Of note, a frame-shift deletion called in the sample RK258 was 52,53 excluded as a false positive after visual inspection in the Integrative Genomics Viewer (IGV). Samples with a structural variation breakpoint on the plus strand 20 kb upstream of *TERT* the transcription start site were defined as TERTp^SV^. TERT^amp^ samples had at least six additional copies of the *TERT* gene compared to the mean ploidy of the sample. Tumor samples with a C228T or C250T *TERT* promoter mutation were defined as TERTp^mut^. Due to the low sequencing coverage at the *TERT* promoter, these mutations were called using less stringent criteria (at least two reads with the mutated base, mutational frequency of at least 20%) in addition to the PCAWG consensus SNV calls (Synapse ID syn7364923). If multiple of these *TERT* modifications were present, the sample was defined as TERT^mult^. Samples with these *TERT* alterations were summarized as TERT^mod^. Samples without any of these alterations were defined as “wild type”. If a sample had both a TERT^mod^ alteration and an *ATRX/DAXX* alteration, it was defined as “mixed”. For some analyses, ATRX/DAXX^non-trunc^, mixed and wild-type samples were summarized as “other”.

### Overlap of juxtaposed positions upstream of *TERT* and predicted super-enhancers

For the closest structural variation (SV) of each tumor sample to the *TERT* TSS, the juxtaposed genomic coordinates were compared to 65,950 predicted super-enhancers from the dbSUPER database^34^. Only SVs on the plus strand and within 1 mb of the *TERT* TSS were considered. Overlaps of juxtaposed positions with super-enhancer sites were defined as direct overlaps. Super-enhancer sites within 1 mb of the juxtaposed position were defined as indirect overlaps.

### Telomere insertion detection

To find insertions of telomeric sequences into non-telomeric regions in the genome, we searched for tumor-specific discordant paired-end reads, where one end was an extracted telomere read and the other end was non-telomeric and uniquely mapped to a chromosome (mapping quality > 30). In 1 kb regions containing at least three discordant reads in the tumor sample and none in the matching control exact positions of telomere insertions were defined by at least three split reads spanning the insertion site. The split reads had to contain at least one TTAGGG repeat. Regions with discordant read pairs in at least 15 control samples were excluded. Finally, the insertion sites were visualized using IGV^52,53^ to identify and remove remaining false positives. A telomere insertion was defined as two-sided if another telomere insertion in opposite orientation was found in the downstream 10 kb of the reference genome. Otherwise it was defined as one-sided.

### Breakpoint detection

Breakpoints were obtained from the consensus breakpoint list of structural and copy number variation calls (Synapse ID syn8042992). In short, six copy number detection tools were run on all samples including the consensus structural variations breakpoints. From the obtained chromosomal segments of the individual callers another set of consensus breakpoints was calculated.

### Chromothripsis detection

To identify chromothripsis events, we extended the set of statistical criteria proposed by Korbel and Campbell^54^. The basic idea is to determine whether there is a statistically significant number of interleaved structural variants (SVs) in a contiguous genomic region. We did this by constructing a graph whose nodes correspond to SVs and whose edges connect interleaved SVs. The identified clusters of SVs were also tested for the presence of alternating copy number and loss-of-heterozygosity patterns. The resulting chromothripsis calls were validated visually. The full description of the methodology and the detailed patterns of chromothripsis events in the genomes will be reported in a separate manuscript by Cortes-Ciriano *et al.* [PCAWG 2017, Cortés-Ciriano *et al.*: Comprehensive analysis of chromothripsis in 2600 human cancers using whole-genome sequencing]. Only high-confidence chromothripsis calls were included in this analysis.

### Copy number changes at telomere insertion sites

Copy numbers of chromosomal segments were obtained from the PCAWG consensus calls (Synapse ID syn8042992). Copy numbers reveal gains or losses of chromosomal segments based on coverage and B-allele frequency, but were here limited to segments of at least 10 kb. The breakpoint estimations could differ from the actual site by up to 50 kb. Therefore, telomere insertions were assigned to the closest breakpoint within 50 kb. If there was no breakpoint within 50 kb or the copy numbers at either side of the telomere insertion were the same, the copy number change at the telomere insertion was defined as neutral.

### Structural variations near telomere insertion sites

Structural variation annotation was obtained from the PCAWG consensus calls (Synapse ID syn7596712), which was based on discordant mate pairs and split reads, providing exact breakpoints. Because copy number variations smaller than 10 kb were not detected by copy number callers, small deletions next to the telomere insertion site may be missed. We therefore searched for structural variations within 10 kb of a telomere insertion to detect these cases.

### Candidate gene selection for correlation analysis

A list of 1,725 telomere maintenance associated human genes was obtained from TelNet^38^ (http://www.cancertelsys.org/telnet/) on February 20 2017. After removing genes without a unique Ensembl IDs in the GENCODE^55^ v19 HAVANA annotation, the remaining 1,686 genes were used for correlation of telomere insertions and simple nucleotide variants.

### Detection of telomere variant repeats

Telomere variant repeats (TVRs) were detected by searching for hexamers of the type NNNGGG in the extracted telomere reads. Each base was required to have a base quality of at least 20. The neighboring 18 bp on either side of the TVR were determined. For further analysis, NNNGGG TVRs were once computed for arbitrary context and once for t-type context ((TTAGGG)_3_-NNNGGG-(TTAGGG)_3_, also called “singletons”). The absolute counts were normalized to the total number of reads in the sample. The expected pattern counts at different telomere content tumor/control log2 ratios were taken from the regression line through TERT^mod^ samples.

### Classifier for predicting active telomere maintenance mechanisms

A random forest classifier to distinguish ATRX/DAXX^trunc^ and TERT^mod^ samples was built using the R packages “randomForest”^56^ and “caret”^57^ with the following eight features: telomere content tumor/control log2 ratio, number of telomere insertions, number of break points and the distance of TGAGGG, TCAGGG, TTGGGG, TTCGGG and TTTGGG singletons (i.e. repeats in a t-type context) to their expected occurrence. To deal with the imbalance in the data set (i.e. 268 TERT^mod^ samples vs. 63 ATRX/DAXX^trunc^ samples without missing data), the model was trained with a down-sampled training set. The performance was determined using 10-fold cross-validation.

### Statistics

Differences between ATRX/DAXX^trunc^ and TERT^mod^ samples in terms of telomere content, percent break points with telomere insertions and singleton repeat abundance were tested using two-sided Wilcoxon rank-sum tests. Singleton repeat abundance p-values were corrected for multiple testing using the Bonferroni method. To reduce the influence of outliers, correlation coefficients were calculated with the Spearman method. Correlation between control telomere content and age as well as tumor and control telomere content was tested with linear regression. All statistical analyses were carried out using R (R Foundation for Statistical Computing).

## Acknowledgments

We thank all PCAWG groups that have provided mutation calls. We thank J. Kerssemakers, M. Prinz and M. Heinold for their help in data processing. We thank I. Buchhalter for annotation of simple nucleotide variations. We thank D. Hübschmann for his support in correlation analysis. We thank all TelNet curators for sharing the results of their extensive literature research. We thank K.I. Deeg, P. Lichter, and S.M. Pfister for their support in early stages of the study. We thank I. Chung for comments and discussion. The work was supported by grants from the German Federal Ministry of Education and Research (BMBF) within the e:Med program (project CancerTelSys, 01ZX1302 to K.R.) and the program for medical genome research (01KU1001A, -B, -C, and -D; 01KU1505A). S.D.K. received funding from the German Research Foundation (DFG) in research priority program SPP1463 (grant no. Br3535/1-2). I.C.C. has received funding from the European Union’s Framework Programme For Research and Innovation Horizon 2020 (2014-2020) under the Marie Sklodowska-Curie Grant Agreement No. 703543.

## Author Contributions

L.S. was involved in all bioinformatical analyses. C.H. performed structural variation annotation, principal component analysis and classification. L.F. was responsible for correlation analysis. S.D.K. analyzed gene expression data and was involved in visualization. L.S., P.G. and L.F. were involved in method development. K.K. and M.S. were involved in copy number analysis. R.K. was involved in data preprocessing. D.M.B. developed and curated the TelNet database. I.C.C., R.X. and P.J.P. provided regions of chromothripsis. B.H. was involved in early stages of experimental design. R.E. was responsible for data management and creating the IT-infrastructure. K.R. and D.T.W.J. provided insights into telomere biology. D.T.W.J. and L.F. conceived the study. B.B. and L.F. oversaw the experimental design and execution. L.S. and L.F. wrote the manuscript with contributions by K.K., D.M.B., M.S., K.R. and D.T.W.J.

---

**Supplementary Figure 1:**
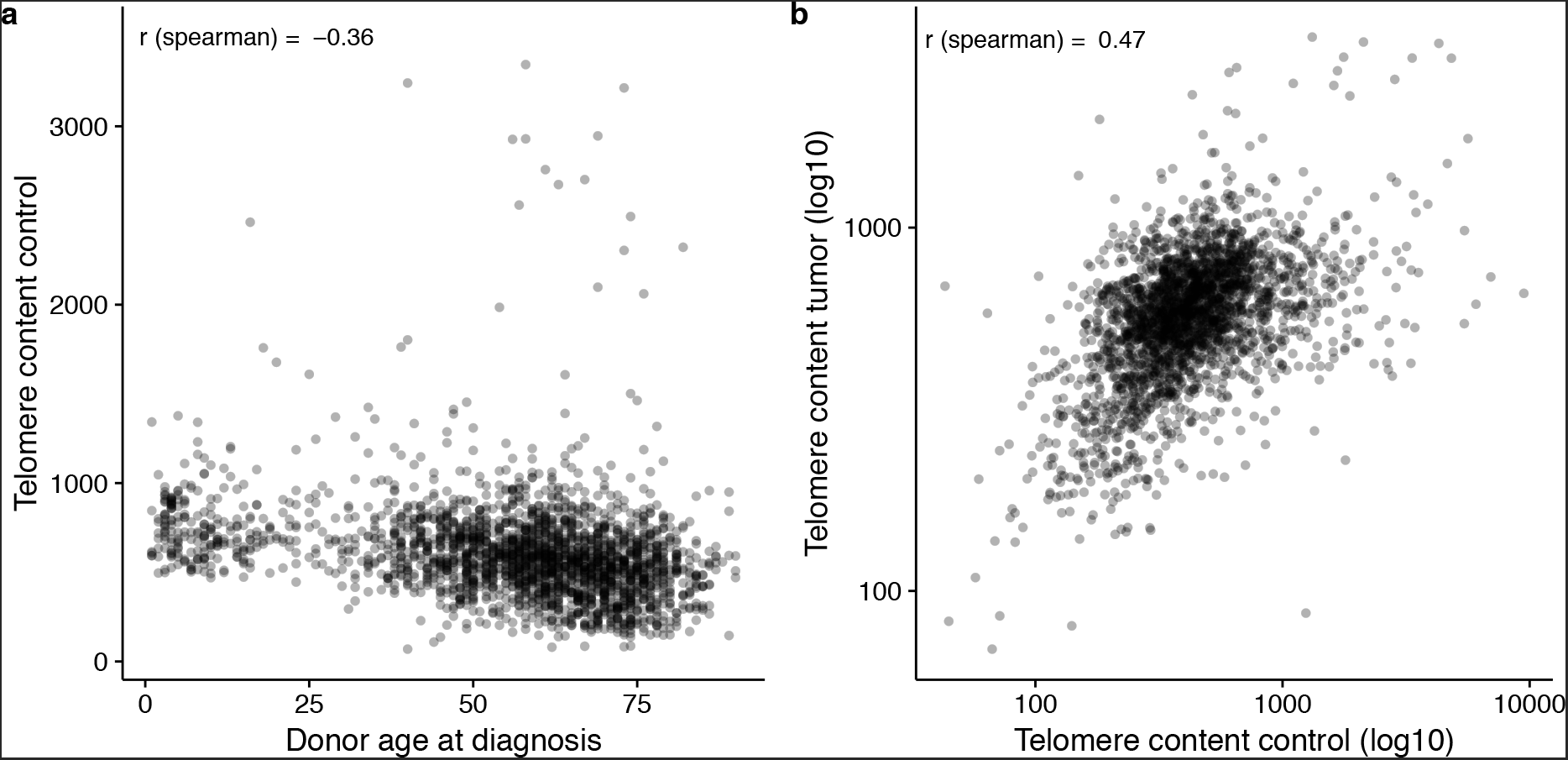
Influences on telomere content. (a) Correlation of control telomere content and the patient age at diagnosis. (b) Correlation of telomere content in the tumor and the control sample.

**Supplementary Figure 2:**
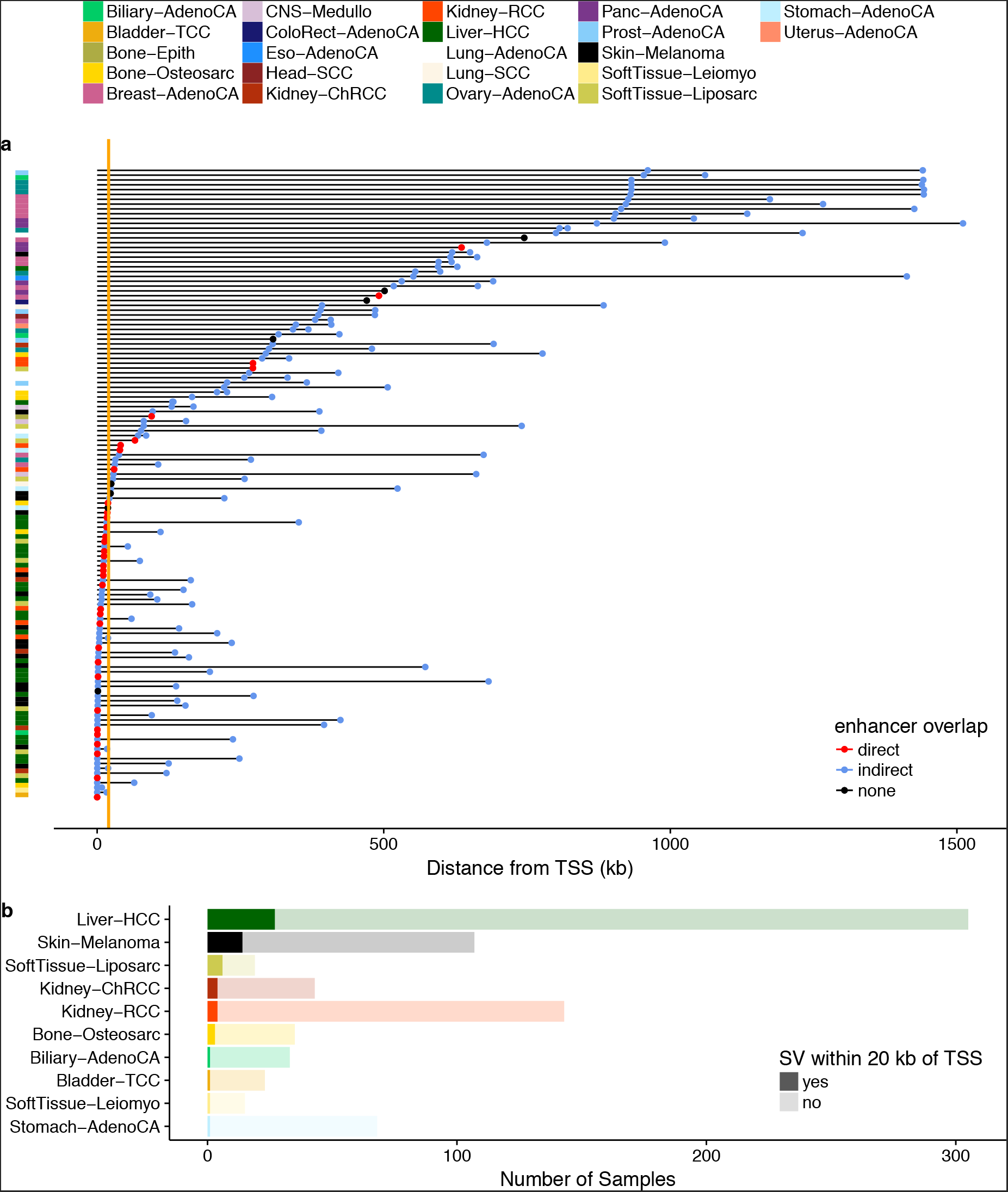
Structural variations upstream of *TERT*. (a) Distance of structural variations (SVs) up to 1 mb upstream of the *TERT* transcription start site (TSS). For each tumor sample, only the SV closest to the TSS is shown. Direct overlaps of juxtaposed positions with dbSUPER enhancer regions are indicated in red. dbSUPER enhancers upstream of the SV are shown in blue, where the first point of each line is the position of the SV and the second point is the rearranged enhancer position. All tumor samples with SVs within 20 kb of the *TERT* TSS (orange line) were considered as TERT^mod^ for the further analysis. (b) Number of samples per tumor type with and without an SV within 20 kb of the *TERT* TSS. Only tumor types with at least one affected sample are shown.

**Supplementary Figure 3:**
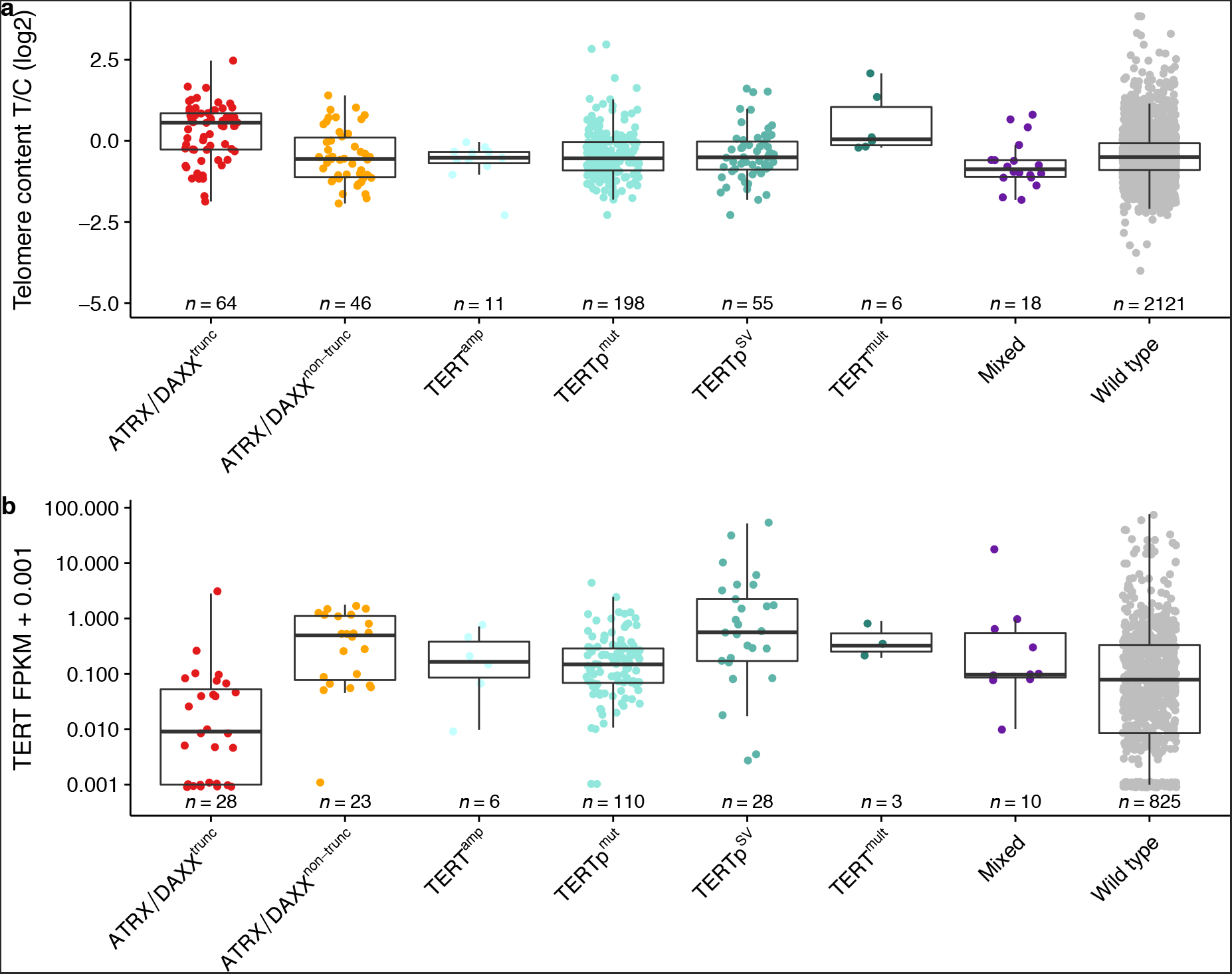
Telomere content and *TERT* expression of tumor samples with different TMM-associated mutations. (a) Telomere content tumor/control log2 ratios. (b) *TERT* expression in FPKMs.

**Supplementary Figure 4:**
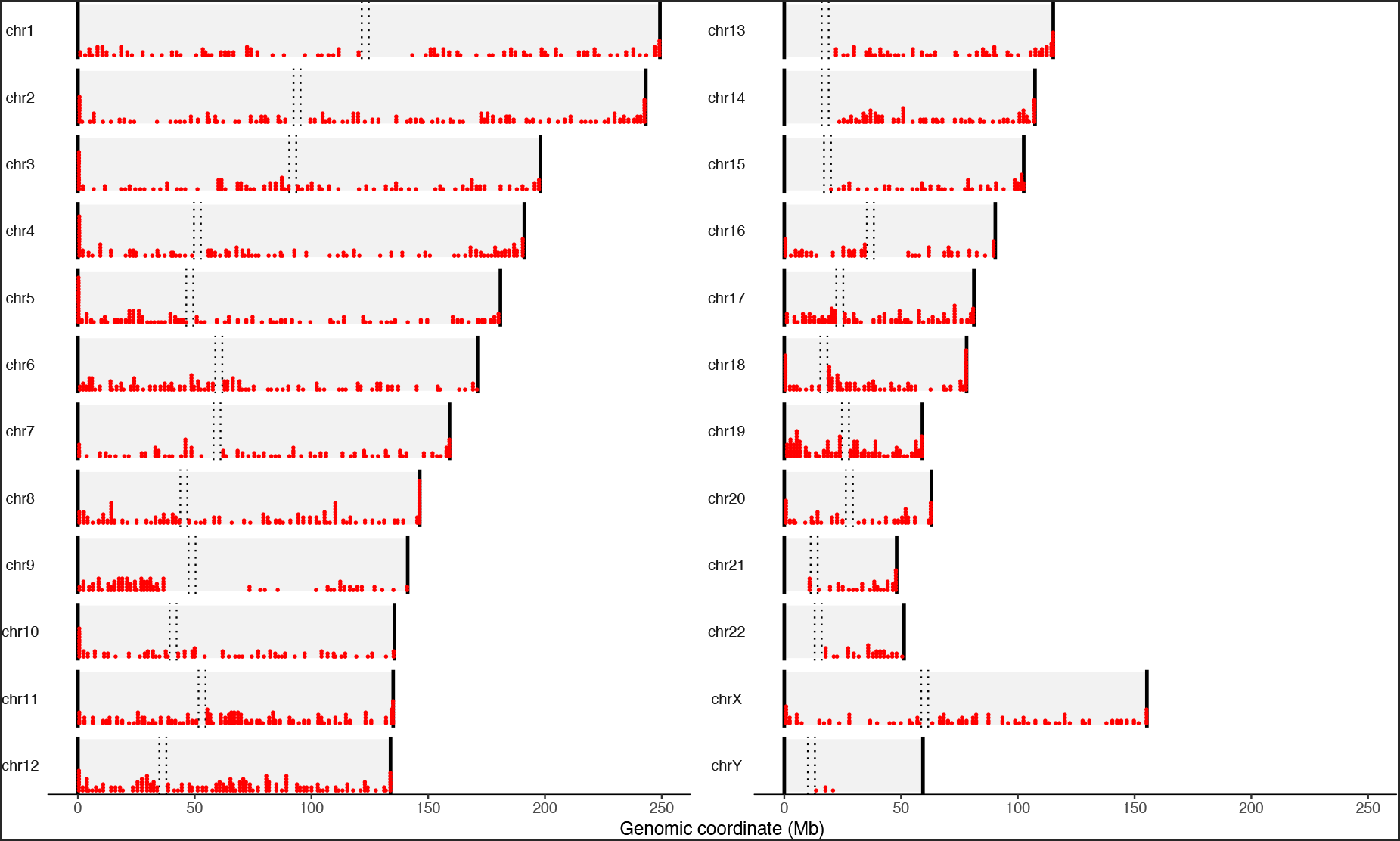
Chromosomal positions of telomere insertions.

**Supplementary Figure 5:**
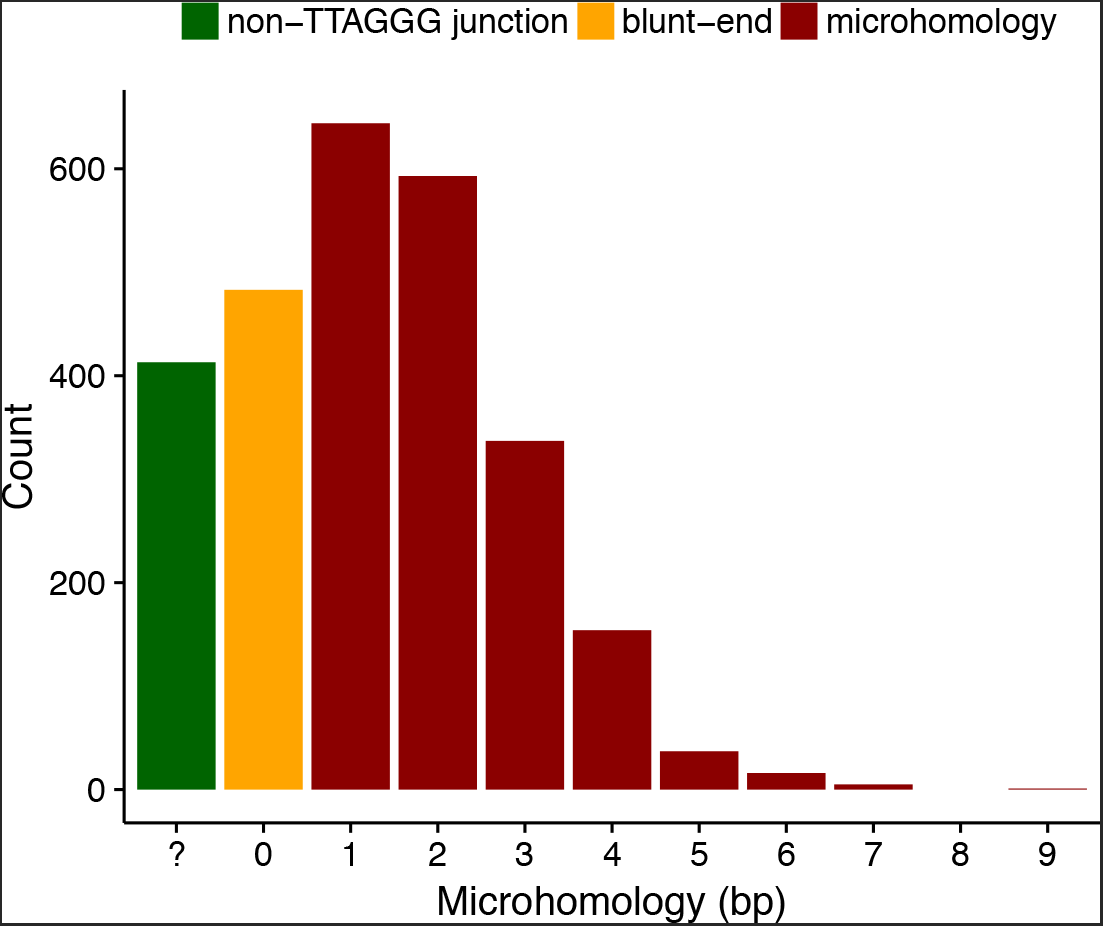
Patterns of microhomology at telomere insertions. The number of homologous bases between the canonical TTAGGG telomere repeat and the human reference genome at telomere insertions is shown on the x-axis. The number of telomere insertions with a pattern of TTAGGG microhomology (red), blunt-end DNA joining (yellow) or without TTAGGG repeats at the junction site (green) are shown on the y-axis.

**Supplementary Figure 6:**
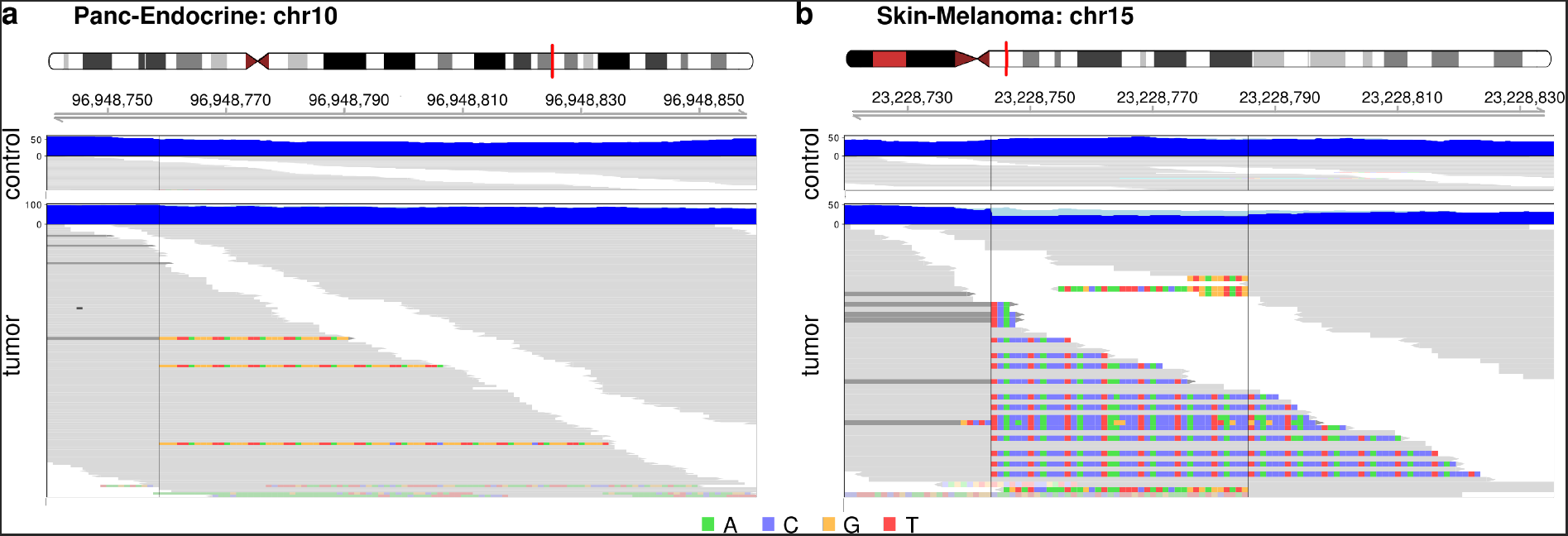
Examples of one-sided telomere insertions without annotated accompanying structural variations. (a) Subclonal telomere insertion in pancreatic endocrine tumor sample SP102547 (copy number at position = 2; tumor purity = 0.87, as determined by copy number calls). Blue tracks show the sequencing coverage; light blue represents clipped sequences. Individual reads are grey and clipped bases are colored. Non-telomeric clipped bases are transparent. Dark grey reads represent the non-telomeric end of a discordant read pair. (b) Unannotated structural variation (position 23,228,744; opaque non-telomeric clipped reads) opposite of a telomere insertion (position 23,228,785) in melanoma sample SP82836.

**Supplementary Figure 7:**
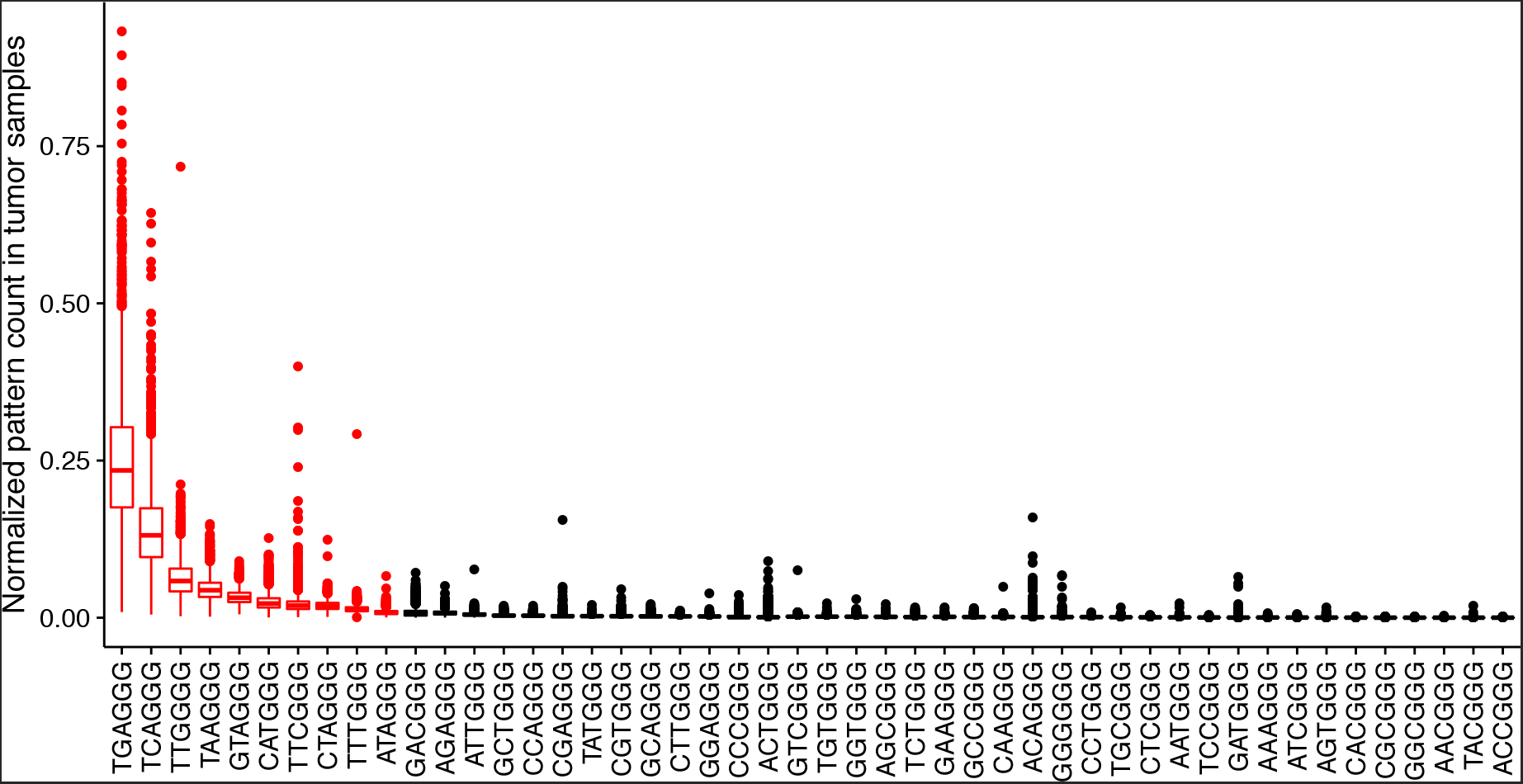
Frequency of TVRs in arbitrary context. The mean pattern counts per telomere read are shown for all tumor samples. TVRs shown in red were investigated further regarding sequence context.

**Supplementary Figure 8:**
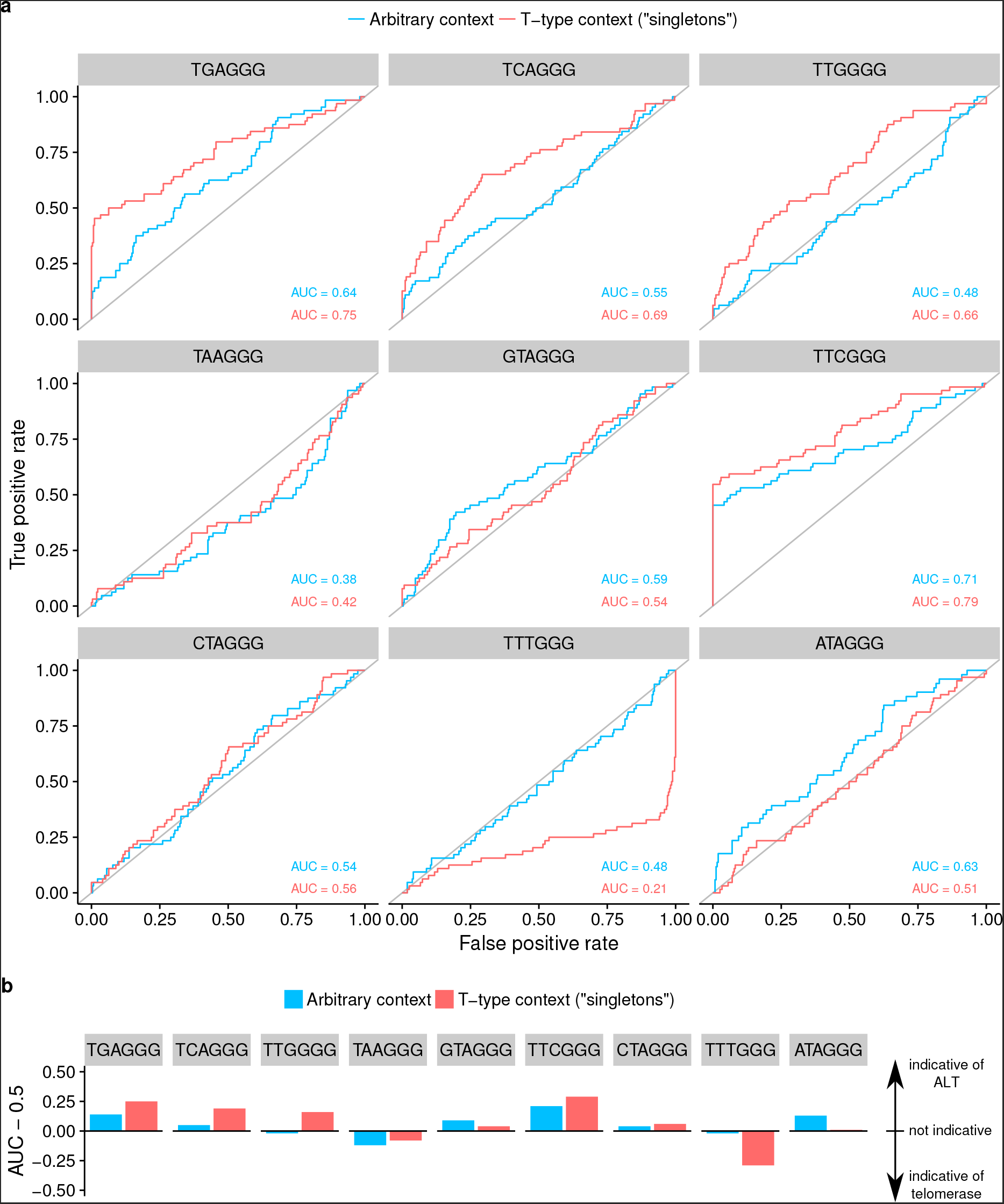
The neighborhood of TVRs is indicative of the telomere maintenance mechanism. (a) Receiver operating characteristic for the classification of samples with ALT-associated mutations from telomere variant repeats. Red: no specific sequence context required. Blue: singletons ((TTAGGG)3-NNNGGG-(TTAGGG)3). (b) Area under the curve (AUC) for the classification of ALT using repeat type counts in different sequence context.

**Supplementary Figure 9:**
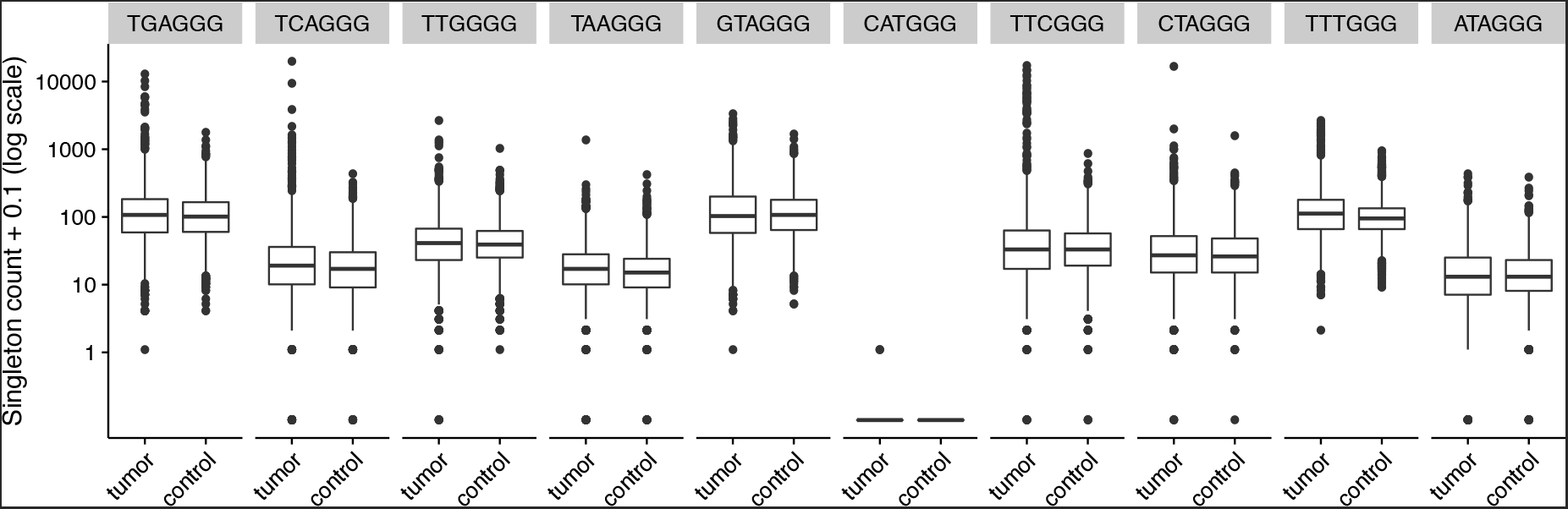
Raw counts of singleton TVRs across all samples.

**Supplementary Figure 10:**
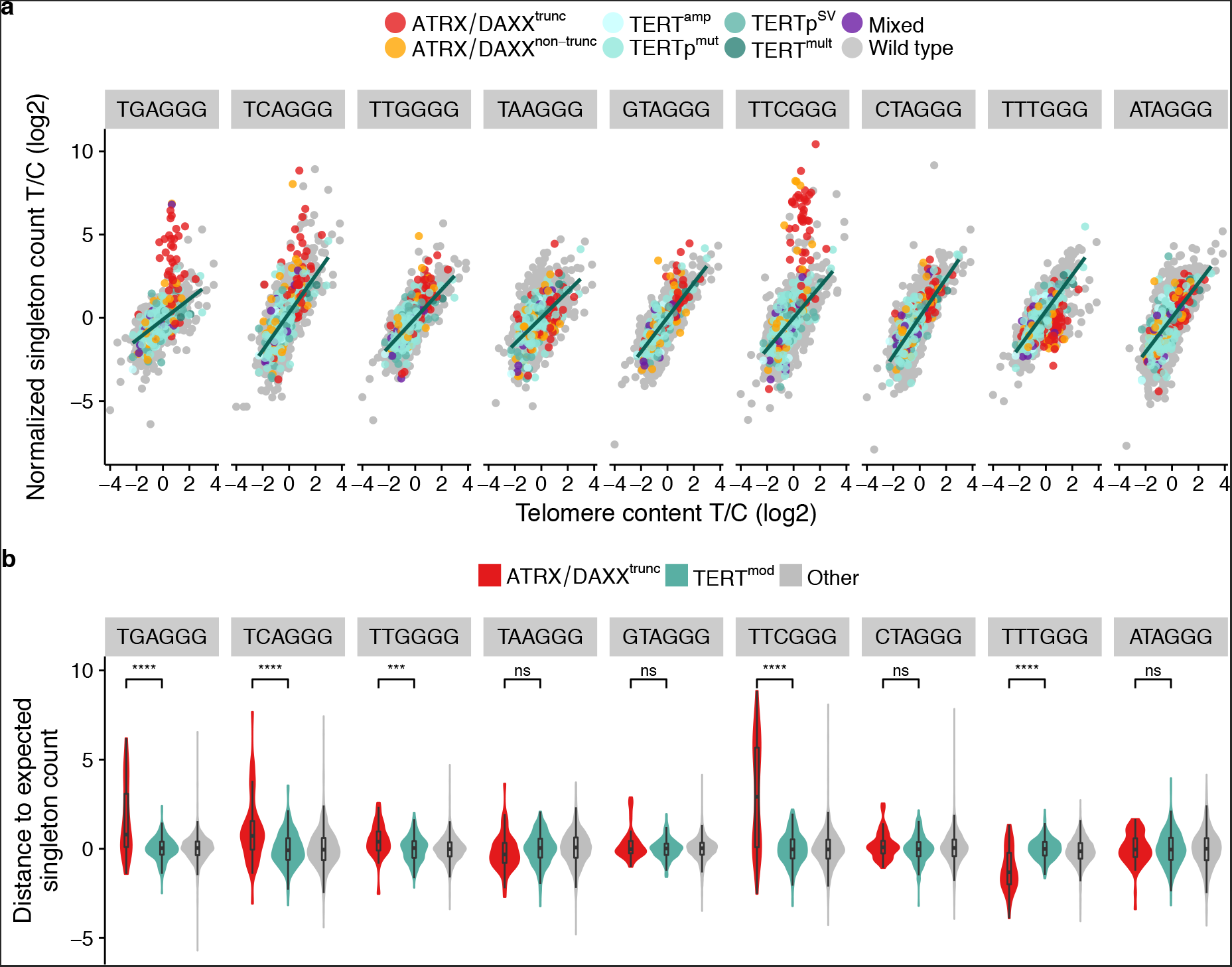
Singleton TVRs. See Figure 4 for details.

**Supplementary Figure 11:**
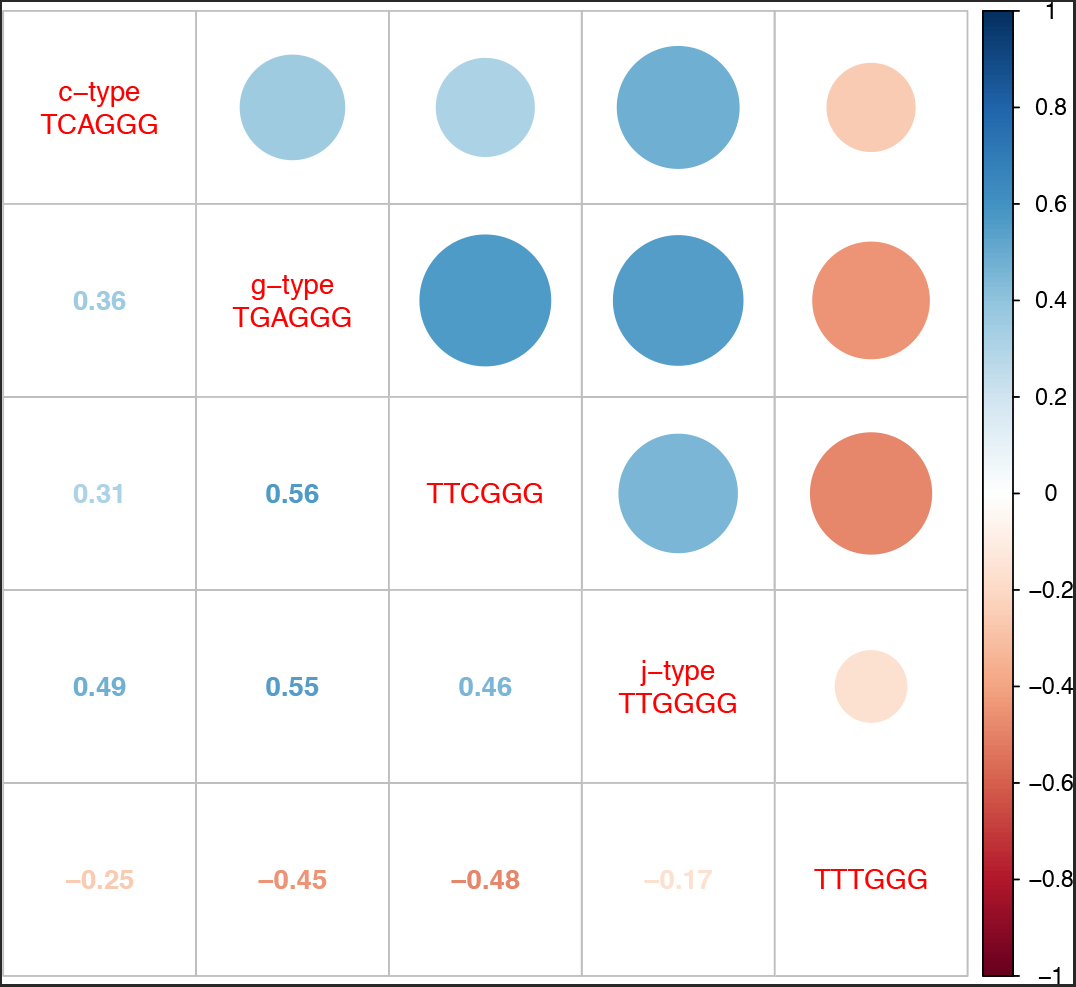
Correlation of singleton TVRs in ATRX/DAXX^trunc^ samples. The Spearman correlation coefficients for the occurrence of the significantly enriched/depleted singleton TVRs in ATRX/DAXX^trunc^ samples is shown.

**Supplementary Figure 12:**
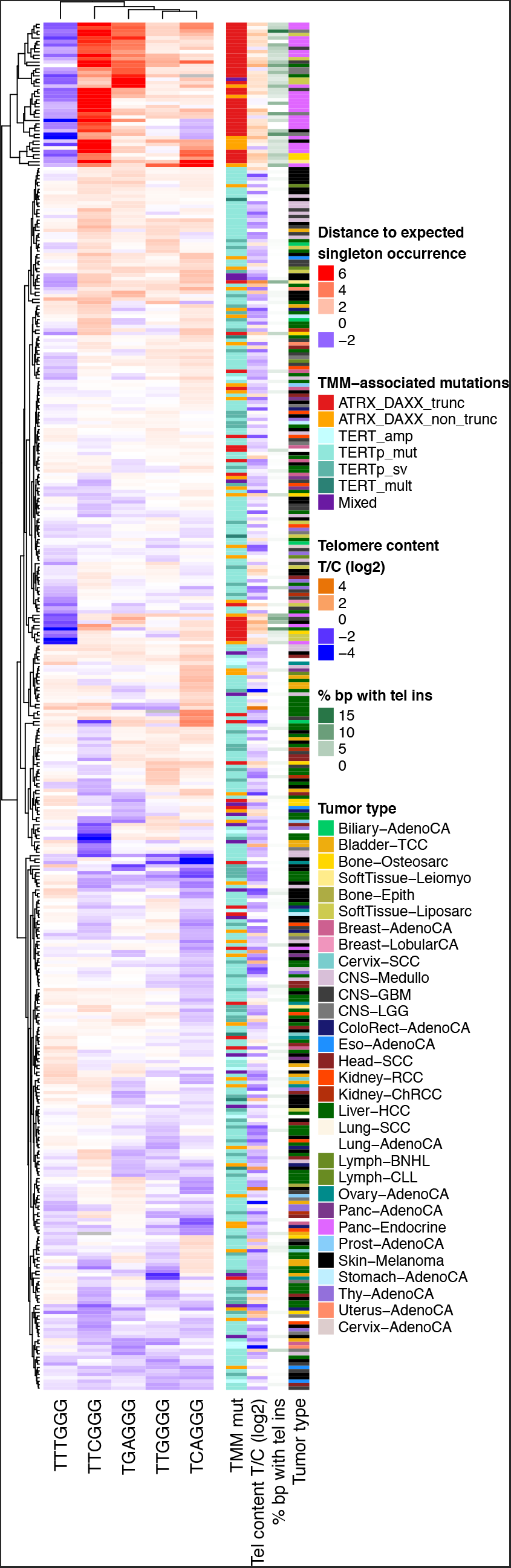
Clustering by singleton TVR occurrences. The heatmap depicts the difference of observed singleton occurrence to the expected occurrence (columns) for tumor samples with TERT^mod^ and/or mutations in *ATRX* or *DAXX* (rows). The TMM-associated mutations, telomere content tumor/control (log2), percent of breakpoints with telomere insertion and tumor type are annotated.

**Supplementary Figure 13:**
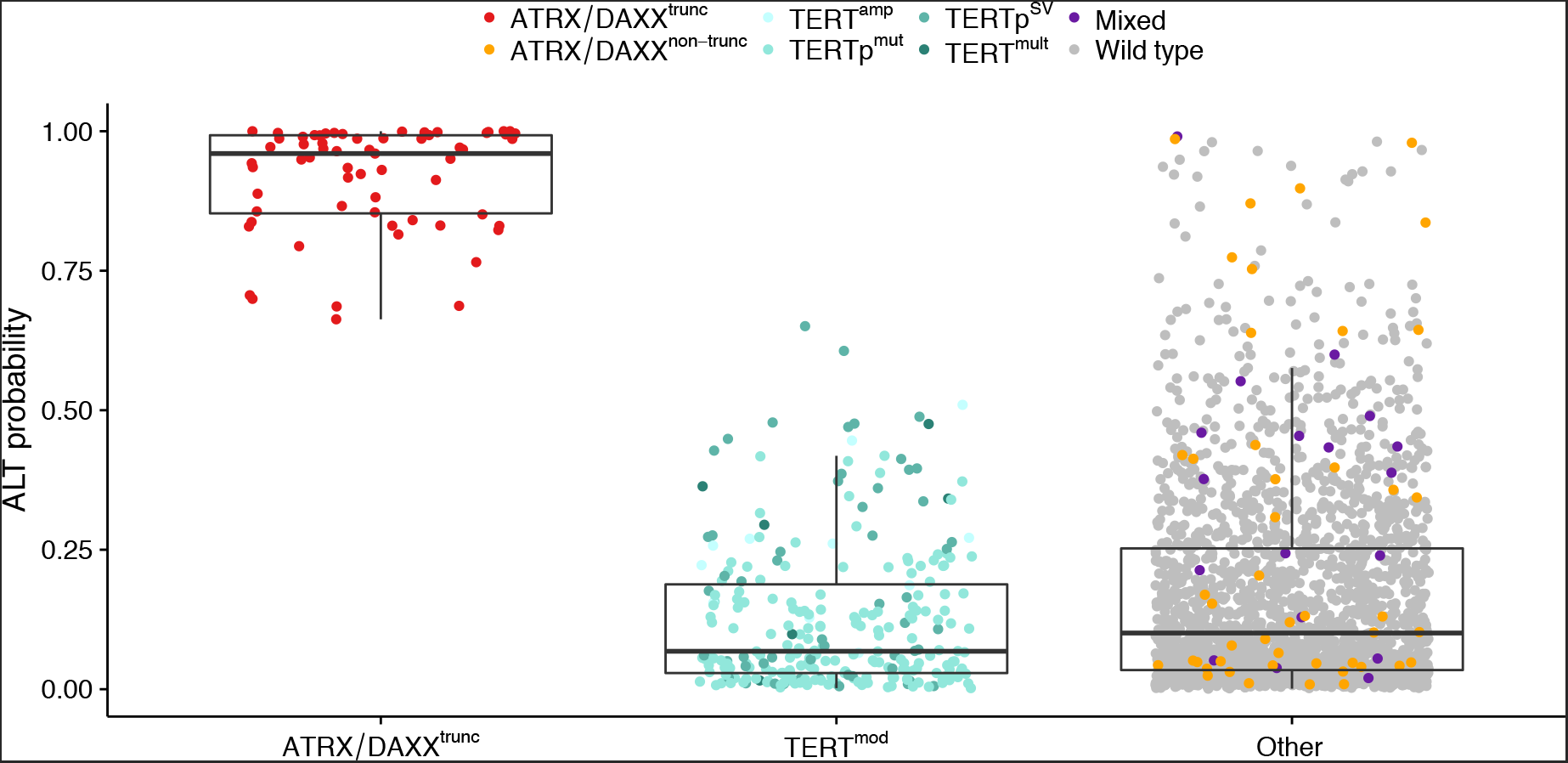
ALT probability of tumor samples with different TMM-associated mutations. The ALT probability was derived from a random forest classifier trained to distinguish ATRX/DAXX^trunc^ from TERT^mod^ samples based on the following features: telomere content tumor/control log2 ratio, number of telomere insertions, number of break points and the distance of TGAGGG, TCAGGG, TTGGGG, TTCGGG and TTTGGG singletons to their expected occurrence.

**Supplementary Figure 14:**
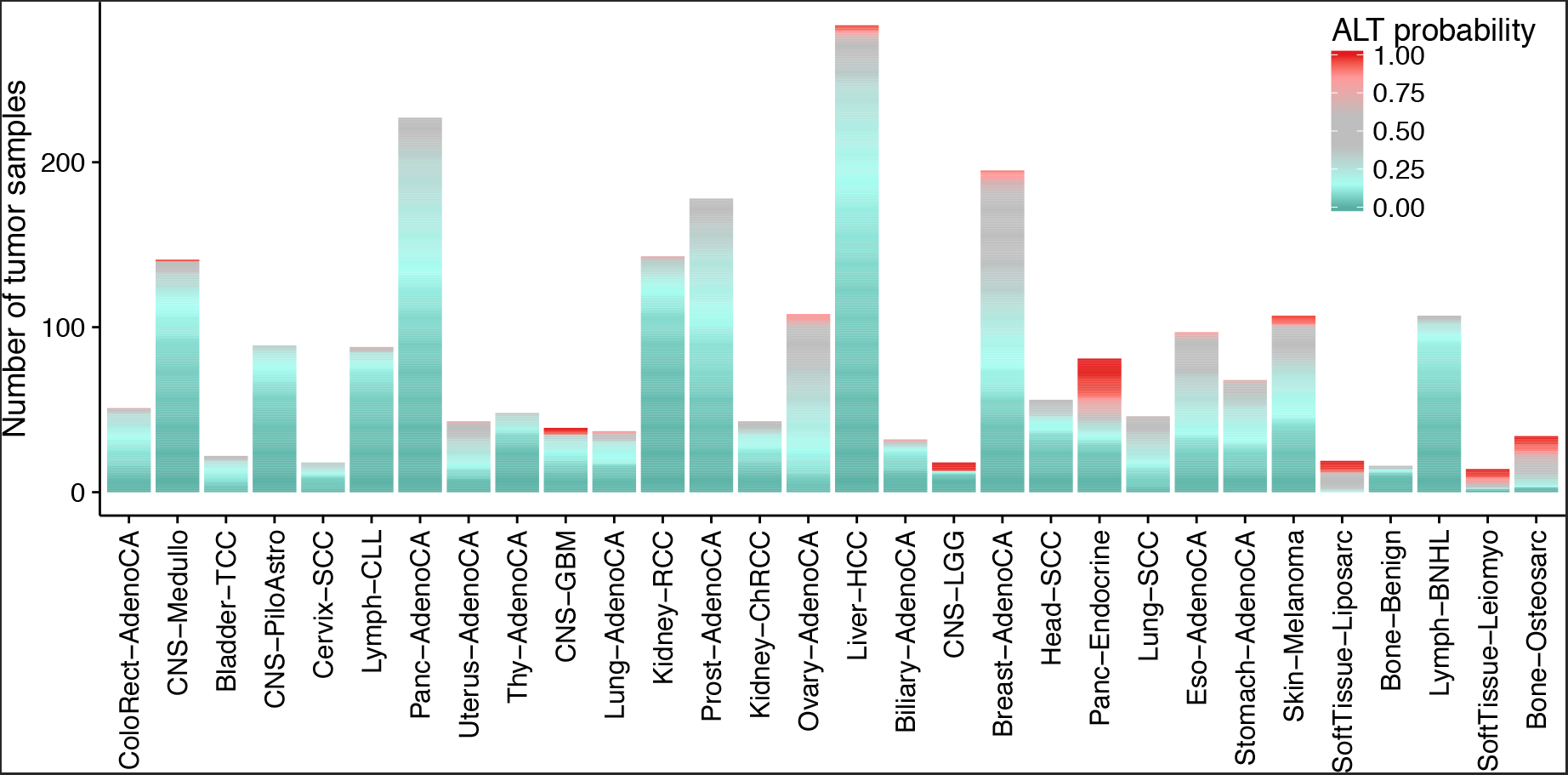
Prediction of ALT probability in different tumor types. For each tumor sample, the ALT probability predicted by a random forest classifier is shown. The tumor types are ordered by mean telomere content tumor/control log2 ratio (from left to right). Cohorts with sample sizes below 15 are not shown.

## ALT classification

The same features that were used for the principal component analysis were used to build a random forest classifier. Using ATRX/DAXX^trunc^ (class I) and TERT^mod^ (class II) samples as the training data set, the classifier achieved an area under the curve of 0.96, a sensitivity of 0.72 and a specificity of 0.98 after 10-fold cross-validation. The variables with the highest importance for the classification were the divergence of observed TTTGGG and TTCGGG singleton TVRs to the expected count, the number of breakpoints and the number of telomere insertions (Supplementary Table 5). The scores resulting from the classifier can be interpreted as an ALT probability. As expected, ATRX/DAXX^trunc^ had a high ALT probability (mean = 0.92), while TERT^mod^ samples had a low ALT probability (mean = 0.13, Supplementary Fig. 13). A total of 18 samples without ATRX/DAXX^trunc^ mutations had a ALT probability of over 0.9, of which two had non-truncating ATRX/DAXX mutations and one sample had a frameshift insertion in *ATRX* and a *TERT* amplification (11 TERT copies, triploid). Across the entire dataset, most samples had a low ALT probability (Supplementary Fig. 14), suggesting that their TMM is telomerase-based. This included some samples with ATRX/DAXX missense mutations, suggesting that the mutations in those samples may be more of a passenger event than functionally relevant. Tumor types with a high ALT-probability were leiomyosarcoma, osteosarcoma, pancreatic endocrine tumors and liposarcomas, in keeping with the known high prevalence of ALT in these entities^58,59^.

